# The splicing factor kinase, SR protein kinase 1 (SRPK1) is essential for late events in the human papillomavirus life cycle

**DOI:** 10.1101/2024.10.28.620581

**Authors:** Arwa A. A. Faizo, Claire Bellward, Hegel R. Hernandez-Lopez, Andrew Stevenson, Quan Gu, Sheila V. Graham

## Abstract

Human papillomaviruses (HPV) infect epithelial causing benign lesions. However, the so-called “high risk” HPVs that infect the anogenital regions and the oropharynx can cause precancerous lesions that can progress to malignant tumours. Understanding the HPV life cycle is important to the discovery of novel antiviral therapies. HPV uses cellular splicing for the production of the full suite of viral mRNAs. Members of the serine/arginine rich (SR) protein family can positively regulate splicing. SR protein activity and cellular location is regulated by kinases through phosphorylation of their serine-arginine domains. SR protein kinase 1 (SRPK1) phosphorylates SR proteins to licence their nuclear entry and can promote nuclear splicing in conjunction with another SR protein kinase, Clk1. SRPIN340 is a specific SRPK1 inhibitor that has been reported to inhibit replication of HCV, Sindbis virus and HIV. Using organotypic raft cultures supporting the HPV16 life cycle, we show that SRPIN340 inhibits the expression of the viral replication and transcription factor, E2. Drug-mediated loss of E2 is specifically associated with inhibition of late gene expression since early gene expression is unaffected. RNA sequencing revealed that SRPIN340 treatment resulted in many gene expression changes opposite to those induced by HPV16 infection. In particular, the loss of epithelial barrier structure and immune function was restored. SRPIN340 treatment resulted in changes in alternative splicing of 935 RNAs and pathway analysis showed a predominance of changes to RNAs encoding proteins involved in chromatin conformation, DNA repair and RNA processing. SRPIN340 treatment was not associated with changes in proliferation or differentiation of keratinocytes. Collectively, these results provide evidence that SRPK1 is a host factor that is essential for HPV16 replication and therefore, targeting this factor, or the phosphorylation events it mediates, could be considered as a therapeutic strategy for HPV16 infection.

**AUTHOR SUMMARY:** Human papillomavirus (HPV) can cause precancerous and cancerous tumours of the anogenital region, for example, the cervix. HPV infects epithelial cells and utilizes the host cell gene expression machinery to produce its own proteins. Splicing is a key mechanism used by HPV to synthesise messenger RNAs that encode the viral proteins. Splicing is positively controlled by SR proteins which require to be phosphorylated by SR protein kinase (SRPK) to enter the nucleus and regulate splicing. SRPIN340 is a specific inhibitor of SRPK. SRPIN340 treatment of HPV16-infected epithelial cells resulted in loss of expression of the viral replication and transcription factor, E2 and abrogation of late events in the viral life cycle. SRPIN340 treatment counteracted some gene expression changes to the epithelium known to be cause by HPV infection. These data suggest that SRPK1 is essential for the HPV life cycle and that it could be a potential anti-viral target.

## INTRODUCTION

Human papillomaviruses (HPVs) are sexually transmitted and infect cutaneous and mucosal epithelia to cause benign lesions or warts [1]. Most HPV infections are asymptomatic and are eventually cleared by the host immune system [2]. However, persistent infection with one of the so-called high-risk HPVs (HR-HPVs), primarily HPV16 and HPV18, can cause lesions in oropharyngeal and anogenital mucosa which may progress to cancer. Globally, HPV16 and HPV18 infections account for up to 70% of cervical cancers, with HPV also linked to cancers in the head and neck, particularly in the oropharynx [3]. Vaccination against the most prevalent HR-HPVs has been delivered over almost the last two decades to young people prior to sexual debut [4]. However, the vaccines are prophylactic which means that there are several decades-worth of infected individuals who have not been vaccinated or have declined vaccination. Therefore, development of antivirals against HR-HPV infection is essential. Understanding viral gene regulation is important to this aim.

The HPV life cycle is intimately linked to differentiation of the epithelium it infects and is tightly regulated by virus-host interactions [5]. HPV early proteins E1 and E2 control viral replication. They are first expressed in basal epithelial cells to establish infection, then are expressed later in the upper epithelial layers to enable vegetative viral genome replication prior to virion formation [6]. E2 is also the viral transcription factor. Viral E6 and E7 proteins, are expressed throughout the life cycle to activate the cell cycle and suppress differentiation of suprabasal epithelial cells while stopping the apoptotic response by degrading p53 [7, 8]. The most abundant late protein is E4, which can alter keratinocyte filaments to facilitate virus replication and egress [9]. The final event in the HPV life cycle is expression of the capsid proteins L1 and L2. Capsid protein expression is restricted to the outermost epithelial layers, an immune-privileged site, to avoid activation of the host immune response [5].

Constitutive and alternative splicing, directed by RNA Polymerase II (RNAPII) transcription and the splicesome, are essential host cell mechanisms for production of multiple mature mRNAs from a single gene. Constitutive and alternative splicing is regulated and enhanced by serine-arginine-rich splicing factors (SRSFs) through their roles in splice site selection, exon definition, spliceosome recruitment and stabilisation [10, 11]. They can bind to exonic and intronic sequence enhancers (ESEs/ISEs) to promote splicing at specific sites. The classical SRSFs include SRSF1-9 which all have a common structure composed of an N-terminal RNA recognition motif (RRM), a C-terminal domain, which functions in spliceosome regulation and protein interactions, and a serine-arginine-rich (RS) domain [11]. The RS domain is subject to phosphorylation by kinases including serine/arginine protein kinase 1 (SRPK1) and CDK-like kinase 1 (Clk1) [12, 13]. This post-translational modification regulates both the function and subcellular localisation of SRSFs [14, 15]. SRPK1 is normally present in the cytoplasm of cells where it phosphorylates newly synthesised SRSFs, to licence their entry into the nucleus [16]. In the nucleus, SRPK1 interacts with Clk1 to promote splicing [13].

HPV gene expression is regulated at the transcriptional and post-transcriptional levels [17]. Production of viral proteins occurs from multiple alternatively spliced mRNAs [18]. HPV utilizes the host cell splicing machinery, including SR proteins, for viral protein expression, but also controls host cell splicing. For example, the E2 transcription factor of the HR-HPVs, HPV16 and HPV31, transcriptionally activates the promoters of the genes encoding SRSFs 1, 2 and 3, up-regulates SRPK1, and can bind SRSFs [19–21].

SRPIN340 is a specific inhibitor of SRPK1 through binding to the enzyme’s active site [22, 23]. SRPIN340 has been shown to have anti-angiogenesis properties by inhibiting VEGF splicing and has been suggested as a potential therapy against exudative age-related macular degeneration and diabetic nephropathy, among other diseases [22, 23]. SRPIN340 has been shown to inhibit replication of HIV, Sindbis virus and hepatitis C virus [24, 25]. SRPK1 and SR proteins are essential factors for the life cycle of other viruses such as herpes simplex virus type 1 (HSV1) [26] and hepatitis B virus (HBV) [27] suggesting that drugs to inhibit SRPK1 could have a general activity against a range of viruses.

We tested the hypothesis that SRPIN340 could inhibit the HPV life cycle through regulating the SR proteins known to be required for HPV mRNA expression [28, 29]. We focused on HPV16 as the most prevalent HPV genotype. We assessed the effect of SRPIN340 on viral protein expression (E2, E4, E6, E7 and L1), and cellular SR proteins (SRSF1-3). We also investigated whether SRPIN340 is associated with toxicity to the host cells in three-dimensional raft cultures. Our results demonstrate that inhibiting SRPK1 using SRPIN340 can inhibit the expression of the HPV E2 replication/transcription factor, and lead to loss of HPV late gene expression. Early gene expression was unaffected. No significant change to epithelial growth or differentiation or to cellular proliferation or viability was detected. RNA sequencing revealed that SRPIN340 induced some changes to keratinocyte differentiation, and to the epithelial barrier, opposite to those caused by HPV infection.

## RESULTS

Work by others has shown inhibition of replication of HIV, Sindbis virus and hepatitis C virus by the SRPK1 inhibitor SRPIN340 [24, 25]. Following on from our previous findings that HPV16 E2 up-regulated SRPK1 during the HPV life cycle [20], we were interested in determining if SRPIN340 could act as an antiviral against HPV. First, we wanted to check if there was any effect of the drug on the growth or morphology of 3D raft tissues supporting HPV infection. We grew raft tissues using HPV16-infected keratinocytes (NIKS16 cells: normal immortalised keratinocytes stably transfected with episomal HPV16 genomes [30, 31] (Supplementary Fig 1)). HPV16 was chosen as the primary focus due to its high prevalence among high-risk HPV types associated with anogenital and oropharyngeal cancers. We chose to test concentrations of the drug at 10 µM or 50 µM since these concentrations had been optimised in previous virus replication studies [24, 25]. Preliminary titration assays confirmed that these concentrations were optimal for effective inhibition of SRPK1 without compromising cell viability in keratinocyte monolayer cultures. Compared to the drug vehicle DMSO, the addition of SRPIN340 in the culture medium after 12 days of tissue growth at either 10 µM or 50 µM, and applied for 48 hours, caused disordered growth of the raft tissues (Supplementary Fig 2A). To minimize the effect of the drug on tissue morphology we developed a method of applying the drug to the upper surface of the tissues in a 50 µl droplet also for the last 48 hours of a 14-day growth period. This volume was chosen because it covered the surface of the tissue without reaching the edge. In contrast to the significant disruption to tissue morphology when SRPIN340 was added to the culture media, there was no observable difference in morphology between DMSO or 10 µM SRPIN340-treated tissues when the drug was applied to the top of the tissues (Supplementary Fig 2B). Some disruption of the upper epithelial layers was observed when the drug was applied at 50 µM (Supplementary Fig 2B).

To test whether SRPIN340 could enter the epithelium upon topical application we applied the fluorophore Alexa 488 in a 50 µl droplet to the top of the raft tissues. This molecule has a similar chemical structure to SRPIN340, which has a molecular weight of 349 Daltons. Alexa Fluor 488 with a molecular weight of 570 Daltons is known to be transmitted between cells by gap junctions, which permit intercellular spread of molecules >1 kDa in size [32]. As a control, topical application of DMSO produced no autofluorescence (Supplementary Fig 2C). Topical application of Alexa 488 revealed penetration down to the basal layers of the tissues and fluorescence was distributed into all the cell layers of the epithelium (Supplementary Fig 2D). This suggests that SRPIN340, which is a smaller molecule than Alexa Fluor 488, could travel through gap junctions throughout the epithelium.

### SRPIN340 treatment leads to reduced phosphorylation and relocation to the cytoplasm of shuttling SR proteins in keratinocytes

To assess whether topically applied SRPIN340 had biochemical effects in HPV16-positive cells, we quantified the levels of SRSF1 and 3, which are phosphorylated by SRPK1 [10], in NIKS16 tissues grown for 12 days and then topically treated for 48 hours with either DMSO or SRPIN340 and compared this with a control SR protein SRSF2, which is a poor substrate of SRPK1. Using a phospho-specific antibody (Mab104) we found that levels of phosphorylated SRSF1 and SRSF3 were reduced upon drug treatment (Fig 1A, C), while levels of SRSF2 did not change significantly, as expected (Fig 1B). Next, we analysed the effect of SRPIN340 on the subcellular location of SRSF1 and SRSF3, whose import into the nucleus is controlled by phosphorylation by SRPK1, and SRSF2, which is mostly confined to the nucleus [10, 11]. Compared to a mainly nuclear location in DMSO-treated NIKS16 cells, SRSF1 and SRSF3 were found in the cytoplasm in SRPIN340-treated NIKS16 cells (Fig 1D, F). In contrast, SRSF2 maintained a nuclear location in both DMSO and SRPIN340-treated cells (Fig 1E). These data confirm efficacy of SRPIN340 in inhibiting the activity of SRPK1 in in HPV16-positive keratinocytes.

**Figure 1.**
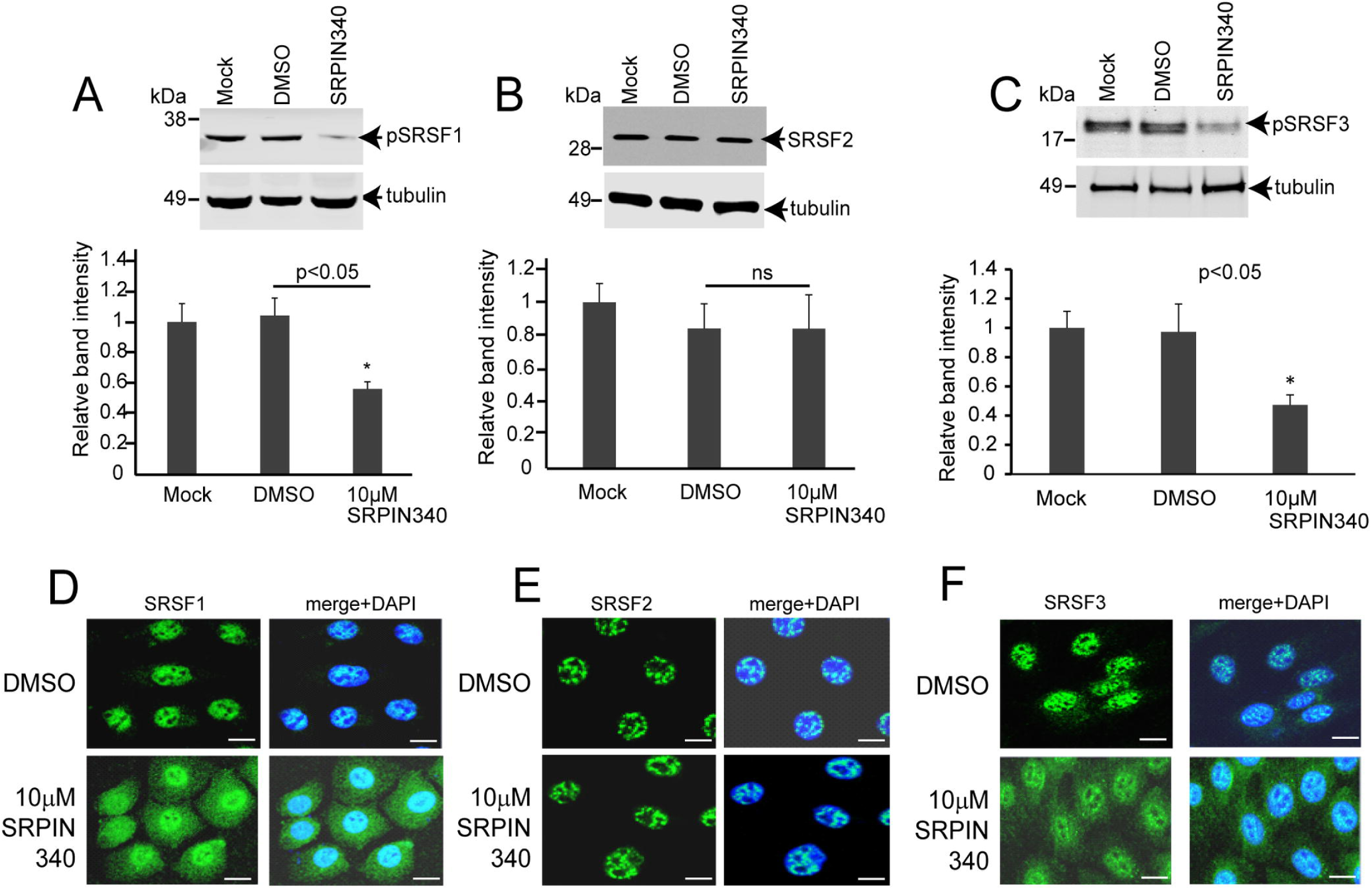
SRPIN340 inhibits phosphorylation of SRSF1 and SRSF3. NIKS16 cells in monolayer culture were mock-treated or treated with 10 µM SRPIN340 or with the drug vehicle, DMSO. A. Representative western blot and graph of the quantification of levels of SRSF1 relative to levels of β-tubulin. B. Representative western blot and graph of the quantification of levels of SRSF2 relative to levels of β-tubulin. C. Representative western blot and graph of the quantification of levels of SRSF3 relative to levels of β-tubulin. The data shown are the mean and standard deviation from the mean from three separate experiments. p<0.05, p-value less than 0.05. *, statistically significant difference in relative protein levels. ns, not statistically significant. D-F. NIKS16 cells treated with DMSO or with 10 µM SRPIN340 D. Immunofluorescence microscopy of cells stained with an antibody against SRSF1. E. Immunofluorescence microscopy of cells stained with an antibody against SRSF2. F. Immunofluorescence microscopy of cells stained with an antibody against SRSF3. The left-hand panels show SRSF staining in green. The right-hand panels show SRSF staining merged with DAPI (blue staining). Representative images from three separate experiments are shown. Size bars = 10µm.

### SRPIN340 treatment does not change E6E7 oncoprotein expression or activity

First, as a measure of HPV early gene expression, we investigated if SRPIN340 as an anti-splicing drug affected the expression of the alternatively spliced mRNAs encoding the HPV16 E6 and E7 oncoproteins (Fig 2A). For these experiments we used NIKS16 cells grown in monolayer culture. End-point PCR analysis showed that the four main splice isoforms of E6E7 mRNAs were present, and their levels were unchanged in the presence of the drug (Fig 2B). Levels of E7 and E6 proteins in NIKS16 cells were much lower than their levels in HeLa cells as a control for oncoprotein detection. Importantly, there was no detectable difference in levels of E6 (Fig 2C, D) or E7 proteins (Fig 2E, F) upon drug treatment. Expression levels of the E6 degradation target protein, p53 and the E7 target, Rb were also unaffected (Fig 2G, H).

**Figure 2.**
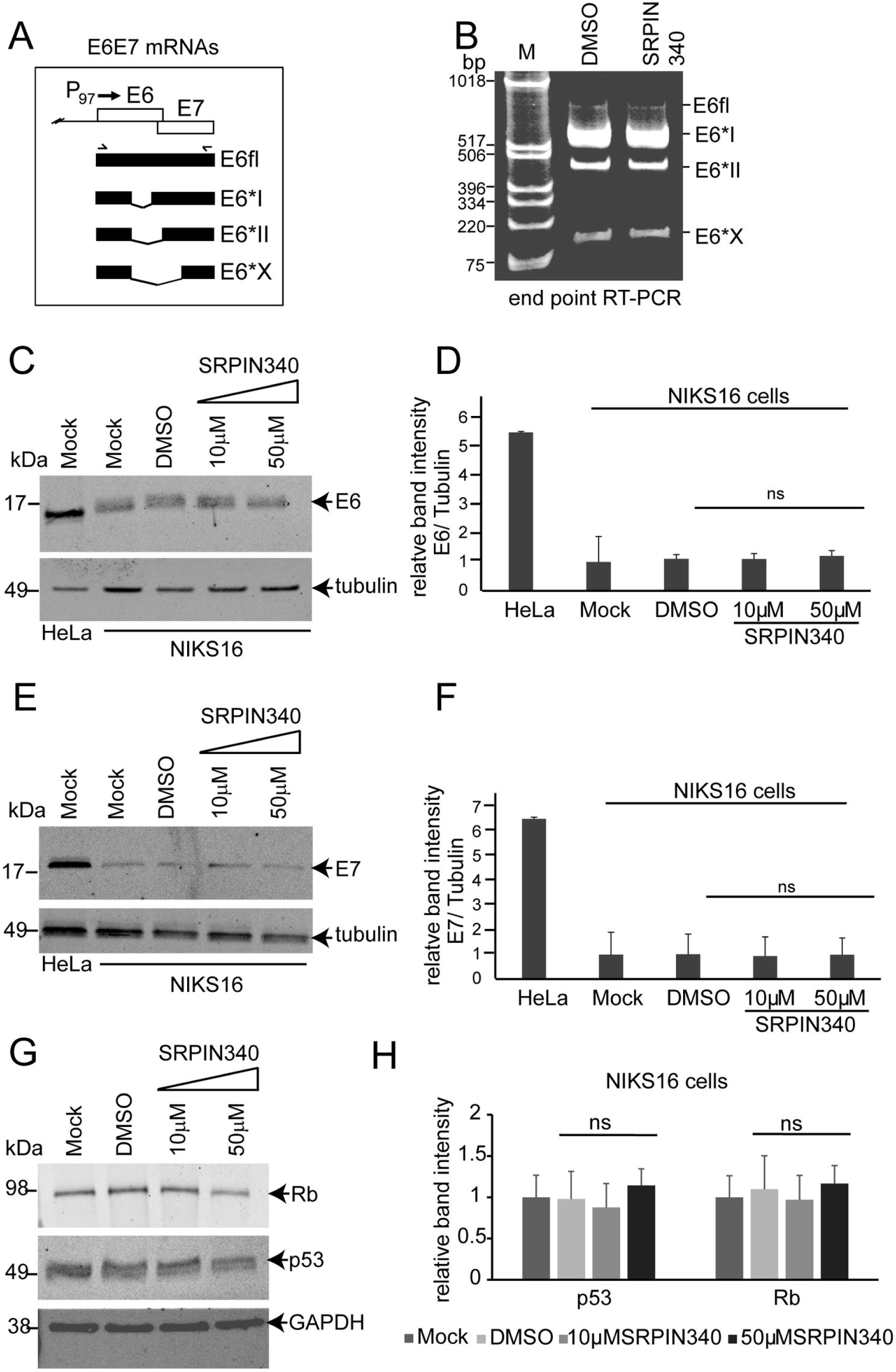
SRPK1 inhibition does not alter HPV16 oncoprotein expression or activity. **A.** Diagram of the E6E7 coding region of the HPV genome showing the unspliced (E6fl) and three spliced RNAs that are transcribed from the region (E6*I, E6*II, E6*X). P_97_, HPV16 early promoter located at nucleotide 97 on the genome. Forward and reverse facing chevrons indicate the approximate position of the primers used in the PCR reaction in (B). B. Gel image showing end point RT-PCR products using the primers indicated in (A) to amplify all four E6E7 mRNAs from NIKS16 cells grown in monolayer culture and treated with DMSO or 10 µM SRPIN340. M = size marker. Western blot of levels of (C) E6 and (E) E7 in monolayer cultured NIKS16 cells either mock-treated or treated with DMSO, or SRPIN340 at 10 µM or 50 µM. HeLa cell extract was included on the blot as a positive control for detection of E6. The upper portion of the blot was reacted with an anti β-tubulin antibody as a loading control. Graph showing quantification of three separate western blot experiments of the levels of (D) E6 and (F) E7 relative to β-tubulin, in HeLa cells as a control and in NIKS16 cells using the various experimental conditions in the western blots. G. Western blots of levels of Rb, p53 and GAPDH as a loading control, in monolayer cultured NIKS16 cells mock-treated or treated with DMSO, or SRPIN340 at 10 µM or 50 µM. H. Graph showing quantification of p53 and Rb levels in the western blot in G. Expression levels were calculated relative to the GAPDH loading control. The data in all the graphs are the mean and standard deviation from the mean from three separate experiments. ns, not statistically significant.

### SRPIN340 inhibits the expression of the viral E2 replication/transcription factor

Next, we investigated the expression of viral late mRNAs using NIKS16 cells grown and differentiated in monolayer culture to express viral late proteins (Supplementary Fig 1B). Expression of the major late HPV16 spliced transcript, E1^E4^L1 and the L2L1 portion of the readthrough transcript E1^E4, E5, L2, L1 (Supplementary Fig 3A) was not altered by SRPIN340 in differentiated NIKS16 cells (Supplementary Fig 3B, C). However, expression of the major mRNA encoding the viral E2 replication/transcription factor [33] (Fig 3A) was inhibited when differentiated NIKS16 cells were treated with the drug (Fig 3B, lane 3). Treatment with DMSO (Fig 3B, lane 1) or with SRPIN349, a SRPIN340 derivative which does not inhibit the kinase activity of SRPK1 [25] (Fig 3B, lane 2), did not affect the expression of the E2 mRNA. Western blot analysis of SRPIN340-treated NIKS16 cells differentiated in monolayer culture revealed that drug treatment inhibited expression of the E2 protein (Fig 3C). Detection of involucrin revealed that the cells were differentiated (see also Supplementary Fig 1B). Quantification of three separate experiments showed a statistically significant difference in E2 levels upon drug treatment (Fig 3D). HPV-negative NIKS cell extract was included in the western blot as a control for the specificity of the E2 antibody. E2 levels were also significantly reduced in drug-treated NIKS16 raft tissues compared to DMSO-treated controls (Fig 3E). Confocal immunofluorescence microscopy confirmed that while E2 expression was detected in the upper layers of NIKS16 3D raft cultures treated with the drug vehicle DMSO, very little E2 was present in drug-treated tissues (Fig 3F). Taken together, these data indicate that SRPIN340 treatment is associated with reduced expression of the alternative spliced HPV16 E2 mRNA, suggesting that SRPK1 kinase activity is important for its expression.

**Figure 3.**
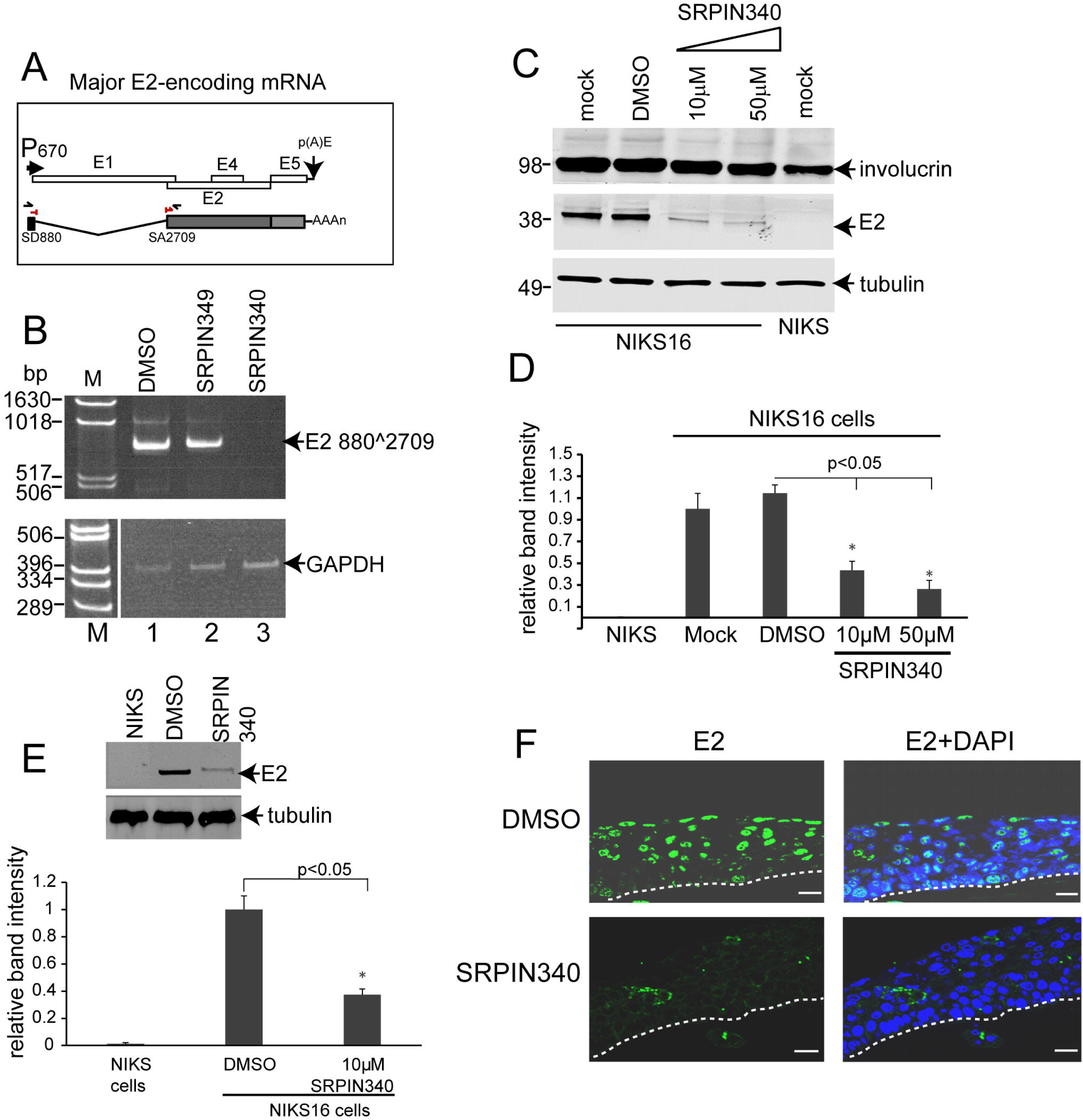
SRPK1 inhibition leads to loss of E2 expression through loss of the spliced mRNA encoding E2. A. Diagram of the early region of the HPV16 genome downstream of the E6E7 gene region. P_670_, HPV16 late promoter located at nucleotide 670 in the E7 coding region. p(A)E, the HPV16 early polyadenylation site. Open rectangles, HPV16 early genes E1, E2, E4, E5. Below is the genome map is shown a diagram of the major spliced mRNA encoding E2. Shaded rectangles, open reading frames. Lines represent RNA sequences which are spliced out to form the E2 mRNA. AAAn, poly(A) tail. Chevrons indicate forward and reverse primers used in the first round of the nested RT-PCR reaction shown in B. A red split chevron indicates the forward primer used in the second round of the nested PCR amplification shown in B. This cross-splice epitope primer was used with the reverse primer located inside the E2 open reading frame. SD880, splice donor site at nucleotide 880. SA2709, splice acceptor site at nucleotide 2709. B. Upper panel: ethidium bromide-stained gel showing the products of an RT-PCR reaction to amplify the E2 mRNA from NIKS16 cells cultured in monolayer and treated with DMSO (lane 1) or with SRPIN349 (lane 2), which does not inhibit SRPK1, or with SRPIN340 (lane 3). Lower panel: GAPDH cDNA was amplified as a loading control. The gel picture is split because the GAPDH reactions were electrophoresed on the right-hand side of the same gel used to visualise the E2 amplification products which were electrophoresed on the left-hand side adjacent to the size markers. C. Western blot showing levels of E2 in monolayer cultured and differentiated NIKS16 cells mock-treated, DMSO-treated or treated with either 10 µM or 50 µM SRPIN340. Protein extract from HPV-negative NIKS cell was included in the experiment to demonstrate specificity of the E2 antibody. The upper portion of the blot was reacted with an anti-involucrin antibody to demonstrate that the cells were differentiated. β-tubulin was used as a loading control. D. Graph showing quantification of E2 levels relative to the β-tubulin loading controls in the various experimental conditions used in the western blot in C. The data shown are the mean and standard deviation from the mean from quantification of three separate western blot experiments. p<0.05*, p-value indicating a significant difference in E2 expression levels. E. Graph showing quantification of E2 levels relative to the β-tubulin loading controls using protein extracts from NIKS16 cells grown in 3D raft cultures. The data shown are the mean and standard deviation from the mean from three separate western blot experiments Representative western blots showing E2 and β-tubulin levels are shown above the graph. p<0.05*, p-value indicating a significant difference in E2 expression levels. F. Immunofluorescence staining with an E2 antibody (green staining, left hand panels) and counterstained with DAPI (blue staining, right hand panels) of sections of 3D NIKS16 raft tissues topically treated with DMSO or with 10 µM SPRIN340. Dotted lines indicate the basal membrane of the tissues. Size bars=50 µm.

### SRPIN340 inhibits late events in the HPV16 life cycle

Loss of the viral replication and transcription factor E2 could result in abrogation of late events in the viral life cycle [6]. We tested this by examining the expression of the major capsid protein L1. Expression of L1 was significantly reduced by SRPIN340 treatment in differentiated NIKS16 cells grown in monolayer culture (Fig 4A, B). Once again, HPV-negative NIKS cell extract was included in the western blot as a control for the specificity of the L1 antibody. Detection of involucrin revealed that the cells were differentiated. A reduction in L1 protein expression was also found in NIKS16 raft tissues grown for 12 days and then topically treated with DMSO or SRPIN340 for 48 hours (Fig 4C, D). E4 is the most abundantly expressed late viral protein. Although antibodies against E4 work poorly in western blot, the E4 B11 antibody can be used successfully in confocal immunofluorescence microscopy. We performed immunofluorescence staining with E4 (Fig 4E) and L1 (Fig 4F) antibodies on sections of DMSO-treated or SRPIN340-treated NIKS16 raft tissues to confirm drug-induced restriction of late events in viral replication. There was a substantial loss of both these late proteins in 3D raft tissues upon drug treatment suggesting that SRPIN340 treatment is associated with inhibition of the late phase of the viral life cycle, potentially due to SRPK1 inhibition.

**Figure 4.**
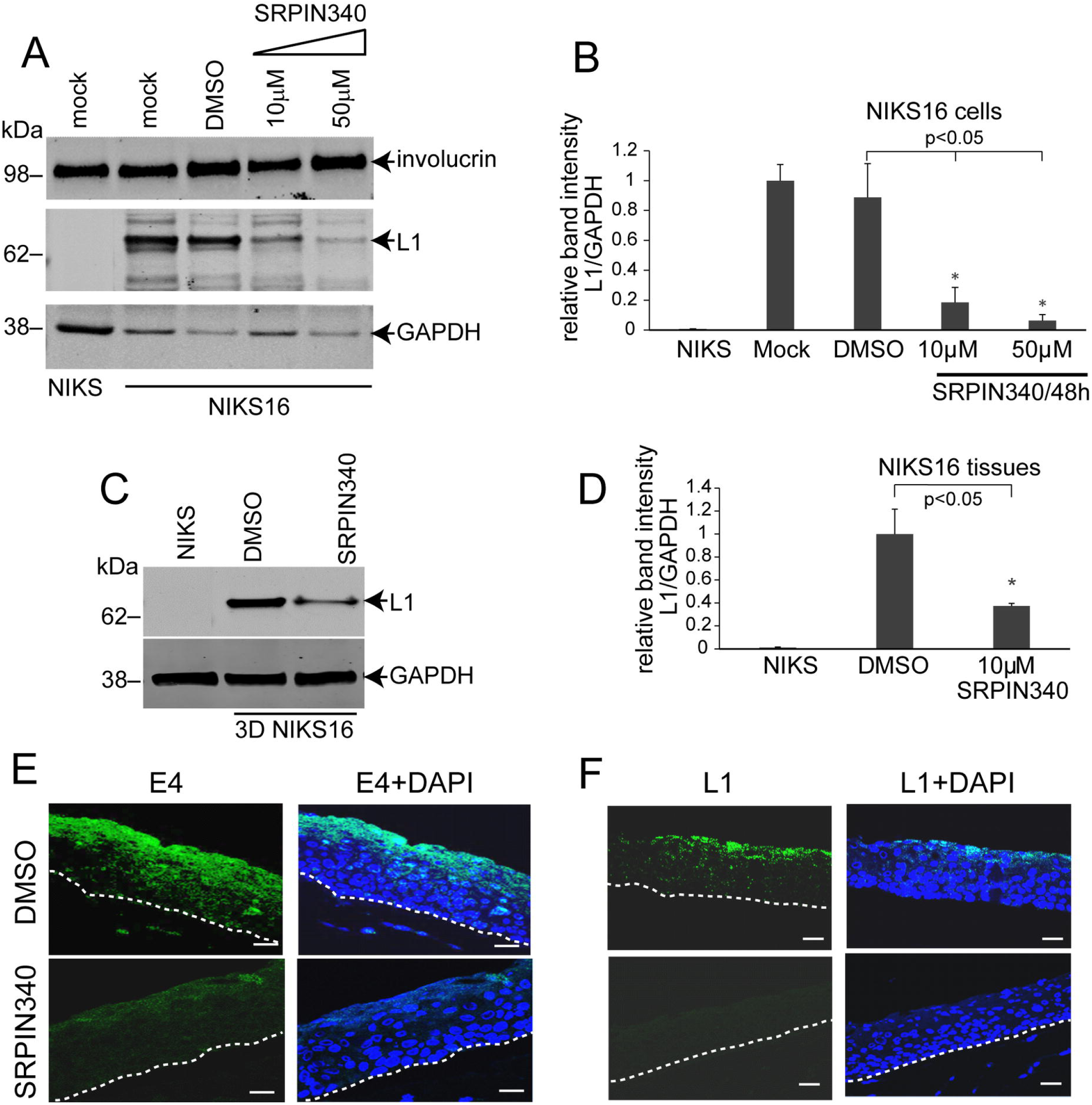
SRPK1 activity is required for HPV16 late gene expression. A. Western blot analysis of levels of L1 protein in monolayer cultured NIKS16 cells mock-treated or DMSO-treated or treated with 10 µM or 50 µM SRPIN340. NIKS cell extract was included on the blot as a control for the specificity of the L1 antibody. The upper portion of the blot was reacted with an antibody against involucrin to demonstrate that the cells were differentiated. GAPDH was used as a loading control. B. Graph showing quantification of L1 levels relative to the GAPDH loading controls in the various experimental conditions used in the western blot in A. The data shown are the mean and standard deviation from the mean from three separate western blot experiments. p<0.05*, p-value indicating a significant difference in L1 expression levels. C. Western blot analysis of levels of L1 protein in 3D raft cultured NIKS16 tissues topically-treated with either DMSO or 10 µM SRPIN340. NIKS tissue extract was included on the blot as a control for the specificity of the L1 antibody. GAPDH was used as a loading control. D. Graph showing quantification of L1 levels relative to the GAPDH loading control in the experimental conditions used in the western blot in C. The data shown are the mean and standard deviation from the mean from three separate western blot experiments. p<0.05*, p-value indicating a significant difference in L1 expression levels. E. Immunofluorescence microscopy of sections of NIKS16 3D raft tissues topically-treated with DMSO or with 10 µM SRPIN340 and stained with an antibody against E4 (green staining, left-hand panels) or counterstained with DAPI (blue staining, right-hand panels). F. Immunofluorescence microscopy of sections of NIKS16 3D rafts tissues treated with DMSO or treated with 10 µM SRPIN340 and stained with an antibody against L1 (green staining, left-hand panels) or counterstained with DAPI (blue staining, right-hand panels). Dotted lines indicate the basal membrane of the tissues. Size bars=50 µm.

### SRPIN340 results in significant changes in gene expression in NIKS16 cells opposite to those induced by HPV16 infection

SRPK1 inhibition can change how SR proteins regulate splicing, leading to changes in the cellular transcriptome. To our knowledge, SRPIN340-induced changes to cellular transcription in keratinocytes have never been reported. In addition, HPV E2 is known to regulate the expression of many cellular genes [34], including SR proteins [19], so loss of E2 expression could contribute to SRPIN340-induced gene expression changes in HPV-infected cells. We carried out RNA sequencing (RNA-Seq) to compare gene expression profiles of SRPIN340-treated to DMSO-treated NIKS16 cells grown in monolayer culture in three replicates of each condition (Fig 5A). Read coverage of the human and viral genomes was not altered by drug treatment (Table 1). However, drug treatment caused a significant change in cellular gene expression (Fig 5A). A total of 342 cellular transcripts were up-regulated, while 241 transcripts were down-regulated (Supplementary Table 1) with a fold change of >log 2.0, Q-value <0.05. Table 2 shows the top 20 up-and down-regulated genes. Ingenuity pathway analysis showed that genes whose expression was up-regulated by SRPIN340 treatment in HPV-infected cells were involved in keratinocyte differentiation, metabolism, immune response and the epithelial barrier (Fig 5B). Down-regulated pathways included aspects of morphogenesis and cell motility, while cell adhesion and the immune system were upregulated, which is consistent with the antiangiogenic activity of the drug (Fig 5C) [22].

**Figure 5.**
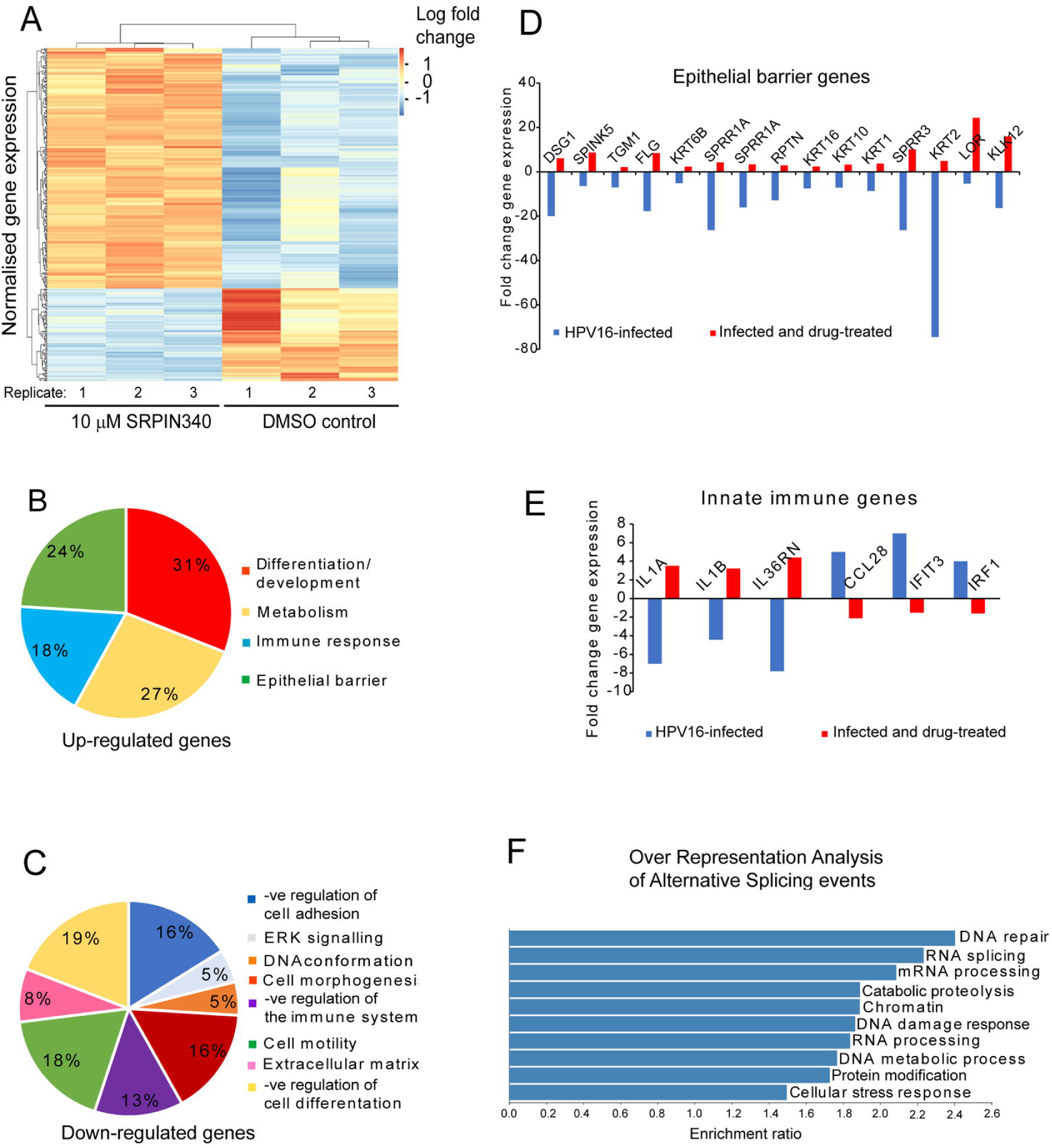
RNA sequencing of SRPIN340-treated tissues reveals that the drug counteracts some of the effects of HPV16 infection on keratinocytes. A. Heatmap of overall changes in gene expression in NIKS16 cells caused by SRPIN340 treatment. Three replicates of DMSO-treated and 10 µM SRPIN340-treated NIKS16 cells were sequenced. B. Pie chart showing the relative percentages of categories of SRPIN340 up-regulated genes in NIKS16 cells. The GO terms for the up-regulated pathways are shown on the right-hand side. C. Pie chart showing the relative percentages of categories of SRPIN340 down-regulated genes in NIKS16 cells. The GO terms for the down-regulated pathways are shown on the right-hand side. D. Graph showing genes involved in the epithelial structural barrier that are down-regulated by HPV16 infection [35] but up-regulated by SRPIN340 treatment. E. Graph showing genes involved in innate immunity that are oppositely regulated by HPV16 infection [35] and SRPIN340 treatment. F. Over representation analysis showing the various cellular processes that are altered by SRPIN340-induced changes in alternative splicing.

**Table 1.** Number of sequencing reads and percent alignment to the HPV16 and human genomes for each sample subjected to RNA sequencing.

|  | Sample | No. of reads | Alignment to HPV16<br>genome (%) | Alignment to Human<br>Genome (%) |
| --- | --- | --- | --- | --- |
| No drug | 1 | 46578063 | 0.04 | 82.10 |
| No drug | 2 | 29538705 | 0.04 | 85.16 |
| No drug | 3 | 37758387 | 0.04 | 80.54 |
| SRPIN340 | 4 | 28468475 | 0.05 | 84.32 |
| SRPIN340 | 5 | 31260092 | 0.05 | 86.22 |
| SRPIN340 | 6 | 32025480 | 0.04 | 80.28 |

**Table 2.** Fold change of the top 20 up- and 20 down-regulated protein-coding genes by SRPIN340 treatment compared to DMSO treatment in NIKS16 cells. Q-value adjusted for multiple hypotheses by DESeq2 analysis.

| Gene | Negative fold<br>change | Positive<br>fold change | Q-value |
| --- | --- | --- | --- |
| TMPRSS11D |  | 6.797647766 | 8.66E-05 |
| SERPINB11 |  | 6.032431007 | 0.000679 |
| CYP1A1 |  | 5.968312781 | 1.21E-25 |
| HRNR |  | 5.415118392 | 0.004903 |
| ENDOU |  | 4.90915265 | 1.9E-06 |
| LOR |  | 4.607461151 | 0.001384 |
| FAIM2 |  | 4.555164059 | 0.010715 |
| LIPN |  | 4.437793821 | 0.017848 |
| SH2D1B |  | 4.393057631 | 2.56E-05 |
| CRNN |  | 4.163890496 | 6.37E-29 |
| CAPN14 |  | 4.084787111 | 1.06E-05 |
| AQP5 |  | 4.069593091 | 6.13E-19 |
| KLK12 |  | 4.00172104 | 4.12E-13 |
| TMPRSS11A |  | 3.98695018 | 3.71E-15 |
| FLG2 |  | 3.887629517 | 9.25E-17 |
| CEACAM5 |  | 3.819780979 | 0.000152 |
| S100A7 |  | 6.797647766 | 8.66E-05 |
| ARG1 |  | 6.032431007 | 0.000679 |
| SPINK7 |  | 3.463702469 | 1.75E-31 |
| SPAG17 |  | 3.4159458 | 8.97E-05 |
| ANKRD1 | -4.307959272 |  | 0.000146 |
| MYL9 | -4.205082349 |  | 6.30E-05 |
| MYL7 | -3.538350233 |  | 2.91E-06 |
| GLP2R | -3.287298644 |  | 0.021162 |
| MGP | -2.883346003 |  | 5.85E-19 |
| KDR | -2.856988802 |  | 3.51E-24 |
| ALDH1L2 | -2.742967901 |  | 0.000597 |
| TGM2 | -2.707864354 |  | 4.79E-17 |
| DRD2 | -2.678088804 |  | 0.007043 |
| ALOX5AP | -2.67413198 |  | 6.89E-06 |
| TAGLN | -2.544919939 |  | 5.02E-34 |
| COL8A1 | -2.329975942 |  | 1.59E-12 |
| SELENOP | -2.313804905 |  | 0.035668 |
| TOX2 | -2.279567504 |  | 0.035591 |
| WFDC2 | -2.742967901 |  | 0.000102 |
| SLC7A7 | -2.087552302 |  | 6.50E-05 |
| ANPEP | -1.964638827 |  | 0.003945 |
| C3 | -1.951676084 |  | 1.11E-30 |
| PTPRB | -1.933491109 |  | 0.00429349 |
| SLCO2A1 | -1.915283426 |  | 1.33E-03 |

We compared the gene expression profiles of SRPIN340-treated NIKS16 cells to HPV16-induced gene expression changes that we identified in a previously acquired dataset comparing compared gene expression changes between uninfected NIKS cells (baseline) and HPV16-infected NIKS16 cells [35]. For this analysis, and to capture the maximum number of genes in common, we focused on genes that exhibited a fold change >log 1.5. Analysis of these changes revealed 91 genes whose expression was altered in both conditions. One important known change to the epithelium caused by HPV16 infection is an abrogation of keratinocyte differentiation and loss of both structural and immune aspects of the epithelial barrier. Comparison of the datasets showed that SRPIN340 treatment led to up-regulation of keratinocyte differentiation genes and epithelial barrier-associated genes that were previously shown to be down-regulated by HPV16 infection. Genes encoding proteins involved in keratinocyte terminal differentiation such as filaggrin and keratin 10, epithelial structural barrier such as loricrin, transglutaminase and repetin [36], and the immune barrier such as keratin 16 [37] and SPRR proteins [38] (Fig 5D), were up-regulated by drug treatment. Moreover, SRPIN340 affected the expression of five genes encoding key immune regulators, including the interferon response factor IRF1 (Fig 5E) which are known to be modulated by HPV16 infection. These findings suggest that SRPIN340 may have the potential to restore some aspects of normal keratinocyte function altered by HPV16 infection.

### SRPIN340 induces alternative splicing changes related to key processes in the HPV life cycle

RNA-Seq data from triplicate 10 µM SRPIN340-treated and DMSO-treated NIKS16 cells were aligned to the human genome and subjected to alternative splicing analysis using the bioinformatic tool SplAdder. ‘Percentage spliced in’ (PSI) values, which are calculated based on the frequency of a particular exon being spliced into a specific splice isoform, were calculated by SplAdder, and used as a measurement of alternative splicing events comparing between SRPIN340-treated and DMSO-treated cells. Differentially spliced genes were then ranked by statistical significance. SRPIN340 treatment resulted in changes in alternative splicing of 935 genes (Supplementary Table 2).

To infer the biological significance of the changes to alternative splicing induced by SRPIN340 treatment in NIKS16 cells, the genes identified as alternatively spliced were subject to pathway analysis. Over-representation analysis was performed on the 935 significantly alternatively spliced genes (p-value <0.05) by Webgestalt [39]. The biological processes affected by the changes in alternative splicing caused by SRPIN340 treatment included RNA splicing, as expected due to the major role of SRPK1 in regulating splicing.

However, splicing of transcripts encoding DNA repair factors/DNA damage response was also significantly altered (Fig 5F). The DNA damage response is essential for HPV replication and alternative splicing-induced changes in DNA repair factors could create a cellular environment inhibitory to HPV replication.

### SRPIN340 is not toxic to keratinocytes

If SRPIN340 can inhibit HPV16 replication it could have therapeutic potential. However, cellular toxicity due to effects on host cell alternative splicing must be considered. We examined the toxicity of SRPIN340 in uninfected NIKS and HPV-infected NIKS16 keratinocytes. Drug concentrations of 10 μM and 50 μM were tested over a period of 72 hours using cells grown in monolayer culture. DMSO was used as the vehicle control. Cell proliferation was not affected by either drug concentration in either cell line (Fig 6A, B). A small decrease in proliferation of NIKS16 cells was observed at the highest drug concentration following treatment for 72 hours, however, this was not statistically significant (Fig 6B). Cell viability of NIKS cells was only decreased significantly at the highest drug concentration following treatment for 72 hours (Fig 6C), while NIKS16 cell viability was unaffected (Fig 6D). Quantification of two cell proliferation markers, Ki67 and MCM2 confirmed that there was no statistically significant change in cellular proliferation upon drug treatment of either NIKS (Fig 6E, F) or NIKS16 cells (Fig 6G, H). These data suggest that short-term SRPIN340 treatment is not toxic to HPV-negative or HPV-positive keratinocytes.

**Figure 6.**
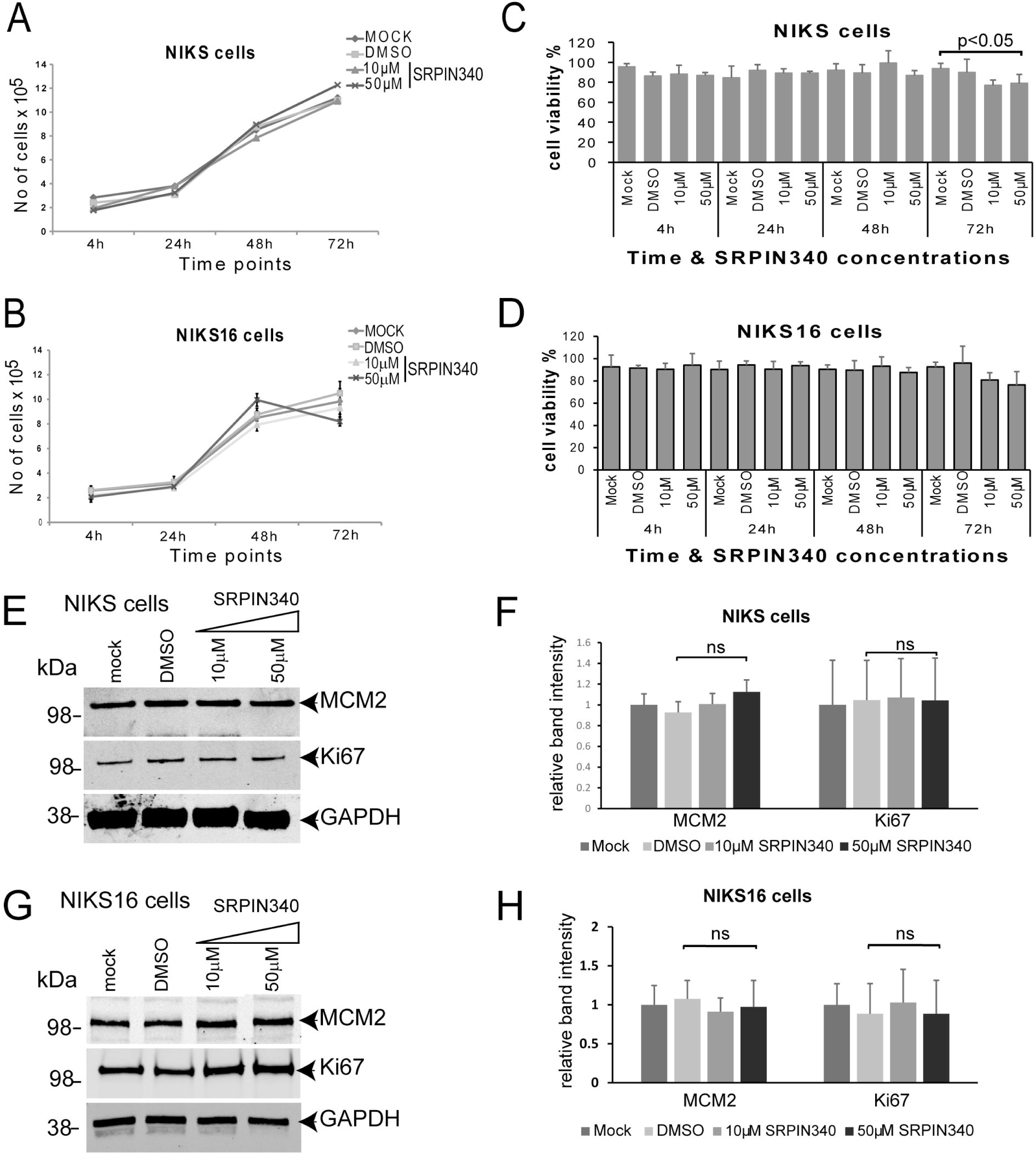
SRPIN340 treatment does not alter cell proliferation or viability in monolayer cultured keratinocytes. A. Proliferation of HPV-negative NIKS cells over 72 hours, mock-treated or following treatment with the drug vehicle DMSO or with 10 µM or 50 µM SRPIN340. B. Proliferation of NIKS16 cells over 72 hours, mock-treated or following treatment with the drug vehicle DMSO or with 10 µM or 50 µM SRPIN340. C. Viability of HPV-negative NIKS cells over 72 hours, mock-treated or following treatment with the drug vehicle DMSO or with 10 µM or 50 µM SRPIN340. D. Viability of NIKS16 cells over 72 hours, mock-treated or following treatment with the drug vehicle DMSO or with 10 µM or 50 µM SRPIN340. p<0.05, p=value of significance. E. Western blot analysis of levels of MCM2 and Ki67 in NIKS cells mock-treated or following treatment with the drug vehicle DMSO or with 10 µM or 50 µM SRPIN340 for 48 hours. GAPDH was used a loading control. F. Graph of quantification of levels of MCM2 and Ki67 relative to GAPDH in western blots of NIKS cell extracts. The data shown are the mean and standard deviation from the mean from three separate western blot experiments. ns, no significant difference. G. Western blot analysis of levels of MCM2 and Ki67 in NIKS16 cells mock-treated or following treatment with the drug vehicle DMSO or with 10 µM or 50 µM SRPIN340 for 48 hours. GAPDH was used a loading control. H. Graph of quantification of levels of MCM2 and Ki67 relative to GAPDH in western blots of NIKS16 cell extracts. The data shown are the mean and standard deviation from the mean from three separate western blot experiments. ns, no significant difference.

## DISCUSSION

Here we have demonstrated that SRPK1 activity is essential for the expression of the HPV16 E2 replication and transcription factor. We showed that key early and late viral mRNAs were synthesised in the presence of SRPIN340. Therefore, transcription of the viral genome is not inhibited by drug treatment, as expected since SRPK1 is associated with post-transcriptional events. Drug treatment did not affect the production of early spliced mRNAs or the late unspliced read-through mRNA containing open reading frames for L2 and L1, suggesting that SRPIN340 specifically inhibits alternative splicing of the E2 mRNA. It is not known which SR proteins are required for E2 splicing. However, a bioinformatic tool, ESE finder [40], predicts with high confidence the binding of SRSF2 and SRSF5 to a sequence 182 bases from the E2 splice acceptor site at nt 2709. Since the activity of SRSF2 is not affected by SRPIN340 treatment, this suggests that E2 alternative splicing is regulated at least in part by SRSF5. Studies by others have shown that hnRNP G has an enhancing effect on splice acceptor 2709 by binding to a sequence between the E2 splice acceptor site and the E2 start codon [41]. Both splicing regulatory factors could cooperate to enhance recognition of the E2 splice site.

E2 regulates alternative splicing of cellular genes [42], which may be due to E2 activating transcription of SR proteins [19]. E2 can also interact with SR proteins and inhibit splicing of suboptimal introns [21]. HPV1 E2 is phosphorylated by SRPK1 [43] but although HPV16 E2 is subject to phosphorylation, it has not been shown to be a substrate of SRPK1, perhaps because it has a short hinge region that does not contain SR repeats. However, the close association of E2 with viral and cellular splicing suggests an important E2-splicing factor regulatory axis in the HPV-infected cell. The requirement for SRPK1 to produce E2 protein and E2 up-regulation of SRPK1 in differentiated epithelial cells [20] underscores this axis and suggests that SRPK1 may be involved in promoting viral DNA replication, genome transcription, and mRNA splicing.

Surprisingly, although SRPIN340 treatment did not inhibit splicing of other major HPV16 mRNAs, it did reduce the cellular levels of HPV major late proteins E4 and L1. This suggests that SRPK1 could positively control viral late mRNA expression at a level other than splicing. SRPK1 is present in the cytoplasm of differentiated HPV16-infected keratinocytes [20]. Therefore SRPK1 could affect mRNA stability, or translation, either directly via its interaction with intracellular signalling pathways [44] or through phosphorylating SR proteins such as SRSF1 involved in these processes [11]. Alternatively, loss of E2 protein, which can activate L1 expression [45] and can bind both E4 and L1 proteins [46, 47], could cause a reduction in levels of both proteins at a post-transcriptional level or perhaps by destabilising the proteins.

Inhibition of late events in the viral life cycle could imbalance the differentiation-dependent life cycle and potentially result in persistent infection and cancer progression. However, we found no evidence for any tumour-promoting activity of SRPIN340 in HPV16-positive keratinocytes, since cell proliferation and viability were unchanged in the presence of the drug, and repression of differentiation due to HPV infection was reversed. Importantly, we found that E6E7 expression and activity were not enhanced by drug treatment.

In our RNA-Seq analyses, SRPIN340 treatment led to up-regulation of expression of keratinocyte differentiation genes with 31% of drug-induced gene expression affecting epithelial differentiation. Keratinocyte differentiation factors included Desmoglein 1, which forms cell-cell junctions to resist mechanical stress and preserve epidermal homeostasis [48], SPINK5 and Kallikrein-12 which are a serine protease inhibitor and serine protease, respectively [49] and loricrin and repetin, which are components of the cornified envelope together with transglutaminase 1, which is involved in its formation [36]. Expression of several keratins was up-regulated as well as filaggrin which binds to keratin fibres [50]. Finally, several of the SPRR proteins involved in epithelial barrier protection and immune defence were also up-regulated [38]. Taken together, the up-regulation of these genes could enhance the epithelial barrier thus counteracting its disruption as a result of HPV16 infection.

This drug-induced effect would be predicted to mechanically strengthen infected, differentiated keratinocytes and inhibit HPV virion egress from the surface of the epithelium. In adddition, we discovered that SRPIN340 treatment of NIKS16 cells changed alternative splicing of RNAs encoding factors involved in cellular processes that are essential for HPV life cycle including DNA damage repair, mRNA processing and chromatin conformation [29]. The effects of SRPIN340 on host cell gene expression may provide an environment in which the host cell can no longer support HPV16 replication, life cycle and transmission.

In monolayer cultures, SRPIN340 had no detectable effect on the proliferation or growth of HPV negative and positive cell lines except when used at high concentration (50 µM). Although there was significant disruption to 3D raft tissues when SRPIN340 was added in the culture medium, there was no change in the morphology or the growth of 3D raft tissues upon topical treatment with the drug at 10 µM. Although animal experiments have demonstrated that systemic oral delivery of SRPIN340 (even at 2000 mg/kg rat body weight) is well tolerated [24] the changes to the epithelium caused by growth of tissues in medium containing the drug suggest that the drug would be best delivered topically if used as an antiviral therapy. SRPIN340 has already been proposed as a topical therapy against diseases such as exudative age-related macular degeneration [22] and next-generation inhibitors to be used at nanomolar concentrations are being developed. VEGF mis-regulation is a key player in HPV-associate cancer progression. SRPIN340 alters VEGF splicing and inhibits angiogenesis [22]. Therefore, although we have studied SRPIN340 activity in the HPV16 life cycle, it may have potential in therapeutic treatment of HPV-associated cancers. Cancer-specific splicing can lead to the expression of oncogenic protein isoforms, as has been shown for HPV-positive HNSCC [51, 52], which SRPIN340 could ameliorate.

There are several limitations to this study. It is important to note that we carried out the analyses in differentiated HPV16-infected, monolayer cultured cells and in differentiated raft cultured tissues so we cannot make any conclusions about the effect of SRPIN340 on early events in the viral life cycle or on persistently infected cells. Additionally, we did not include uninfected keratinocytes treated with SRPIN340 in our transcriptome analyses. Therefore, we cannot conclusively determine whether the observed gene expression and splicing changes upon SRPIN340 treatment are specific to HPV16-infected cells or represent general effects on keratinocytes. Moreover, it would be important in future to determine if SRPIN340 could alter expression of E2 from other high-risk HPVs, or from low-risk genotypes that cause genital warts, or even those HPVs such as HPV1 that cause common warts. The long-term effects of SRPK1 inhibition on epithelial cell function warrant exploration, as sustained inhibition could impact processes such as cellular differentiation and tissue repair. Understanding these effects will be essential for developing safe and effective therapies targeting SRPK. Given that SRPIN340 targets a host factor, combining it with direct-acting antivirals could yield synergistic effects, enhancing antiviral efficacy while potentially lowering the risk of resistance.

## CONCLUSIONS

We have identified host cell gene expression changes following SRPIN340 treatment in HPV16 infected cells. The anti-viral effect of SRPIN340 may be a combination of down-regulation of production of the viral replication and transcription factor E2, together with changes to splicing of host cell transcripts which collectively may alter the host cell environment such that it can no longer support late events in the HPV16 life cycle. The inhibition of HPV16 replication by SRPIN340 suggests that SRPK1 is a host factor crucial to the HPV life cycle.

## MATERIALS AND METHODS

### Cell Culture

NIKS16 cells are normal spontaneously immortalised keratinocytes which have been transfected with HPV16 genomes [30, 31]. NIKS16 cells were grown in monolayer culture on a feeder layer of mitomycin-C-treated J2 3T3 cells as previously described [53] and were used at a maximum passage number of 15 to avoid genome integration. NIKS16 were induced to differentiate by growth at a high density in the presence of 1.2mM Ca^2+^ [19]. Cells were maintained in 5% CO_2_ at 37°C. NIKS16 cells were treated with either DMSO or SRPIN340 (Sigma, UK: SML1088) dissolved in DMSO at the given concentrations on day 8 post-seeding into culture plates and cells were incubated in the presence of the drug for 48 hours.

Raft cultures were established as previously described [54] by seeding NIKS16 cells onto the top of collagen matrices in 24 well plates and incubated at 37°C in 5% CO_2_ for 2 days in E-medium until a monolayer was formed. Rafts were lifted to the air-liquid interface on a sterile stainless-steel wire grid and cultured for 14 days. The E-medium was changed every other day. DMSO or SRPIN340 (Sigma) dissolved in DMSO was added to the culture medium or placed on top of the raft tissues in a 50 µl droplet at the described concentrations on day 12 following lifting to the air-liquid interface and incubated for 48 hours.

### Cell growth kinetics, viability and tissue morphology

Formalin fixed tissue cultures were paraffin-embedded and sectioned (2.5 μm sections). Sections were haematoxylin and eosin-stained and morphology was examined under an Olympus Bx57 light microscope. For cell viability and estimation of cell number, cells were harvested by trypsinization, diluted in 0.1% trypan blue at a ratio of 1:1, and counted using a haemocytometer. The number of unstained (viable) and total number of cells were determined.

### Tissue permeability of Alexa fluor 488

Cell to cell gap junction connectivity and raft permeability were assessed by the ability of cells to transfer the Alexa Fluor 488 dye (A10436, Invitrogen). Alexa Fluor 488 dye at 10 µM was added to the top of 3D cultured tissues in a 50 µl droplet on day 12 of tissue growth and allowed to spread through the tissue structure for 2 days. At day 14, tissues were fixed, sectioned, counterstained with DAPI and dye spread between neighbouring cells from the top to the bottom of the raft structure was visualized using a Zeiss LSM 710 confocal microscope. Zen black software (Zeiss) was used for capturing images.

### Southern blotting

DNA was isolated using the QiaAmp DNA mini kit (Qiagen, Germany) according to the manufacturer’s instructions then subjected to restriction enzyme digestion overnight at 37°C with 25 units of enzyme per 20 µg DNA. To prepare the DNA probe, plasmid pEF-HPV16 was linearised by digestion with Bam HI at 10 units per µg DNA for one hour at 37°C. The digestion products were fractionated on an agarose gel and the 8 kb viral genome band was excised and purified using the QIAquick gel extraction kit (Qiagen, Germany) according to the manufacturer’s instructions. DNA was denatured by boiling for 5 min followed by chilling on ice. The denatured DNA was labelled with Biotin-16-dUTP (Roche #11093070910, Merck, UK) using the random primer labelling system according to the manufacturer’s protocol (Roche #11004760001, Merck, UK). Restriction enzyme-digested NIKS16 DNAs and, as a size control, pEF-HPV16 digested with Bam HI, were fractionated on a 0.8% agarose gel in 1 x Tris borate electrophoresis buffer overnight at 20 volts. Gel treatment, DNA transfer onto nitrocellulose membrane and hybridisation with the pEF-HPV16 probe was carried out exactly as described in the Amersham Hybond N+ protocol (Cytiva, UK).

### Western blotting

Cells in monolayer were scraped into NP-40 lysis buffer (50 mM Tris-HCl pH 8.0, 150 mM NaCl, 0.5% (v/v) NP-40 (IGPAL CA-630)), containing PhosphoSTOP (Merck, UK catalogue # 04693116001) and complete miniprotease inhibitor cocktail (Merck, UK, catalogue # 200-664-3). Protein extraction from raft culture was carried out by adding a small amount of liquid nitrogen to the tissue and grinding to a powder in a mortar and pestle. NP-40 lysis buffer containing PhosphoSTOP (Roche) and complete miniprotease inhibitor cocktail as above was added to the ground tissue, mixed and collected in 0.5ml Eppendorf tubes. The insoluble matrix was removed by centrifugation for 20 min at 13,000 g at 4°C. All protein concentrations were determined using Bradford’s assay and a NanodropTM spectrophotometer (Thermo Scientific).

Protein samples were resolved by 4-12% Bis-Tris SDS-PAGE (Invitrogen, UK catalogue #NP0322BOX) using 1X NuPAGE MES gel running buffer (Invitrogen, UK, catalogue # NP0002). Separated proteins in the gel were electro-transferred onto nitrocellulose membrane at room temperature using the iBlot™ Dry Blotting System (Invitrogen, UK). Membranes were blocked in 5% foetal bovine serum (FBS) in phosphate buffered saline (PBS) for 60 min at room temperature. Membranes were incubated with primary antibody diluted in PBS-T (PBS with 0.1% Tween) containing 5% FBS overnight at 4°C before being washed three times with PBS-T. Primary antibodies were SRSF1 (1:1000, Mab96, Thermo Fisher Scientific, catalogue # 32-4500), SRSF2 (1:1000, Abcam, catalogue #Ab204916), SRSF3 (1:500, Life Technologies, UK, catalogue #334200), SRPK1 (1:500, clone G211-637 BD Transduction Laboratories, catalogue #611072), α-tubulin (1:5000, clone A11126, Thermo Fisher Scientific catalogue # 236-10501), involucrin (1:1000 clone SY5, Merck, UK, catalogue #19018), GAPDH (1:1000, Meridian Life Sciences, UK, catalogue #H86504M, clone 6C5), HPV16 E2 (1:500 Santa Cruz, Germany, catalogue #sc-53327, TVG271), HPV16 E6 (1:500, Santa Cruz, Germany, catalogue # sc-365089), HPV16 E7 (1:500 Santa Cruz, Germany, catalogue #sc-51951), HPV16 L1 (1:500 Dako, Denmark, clone K1H8), p53 (1:700, BD Pharmingen, UK, catalogue #554294), Rb (1:500, Cell Signaling Technologies, Germany, catalogue #9309, clone 4H1), MCM2 (1:500, Abcam, UK, catalogue #Ab133325), Ki67 (1:1000, Abcam, UK, catalogue #Ab197234), keratin 10 (1:500, Abcam, UK, catalogue #Ab9025). Monoclonal antibody104 (Mab104), which detects phosphorylated SR proteins, was prepared from hybridoma supernatants (ATCC CRL-2067) and used neat. Secondary antibodies were diluted in PBS-T containing 5% FBS and incubated with the membrane for 1h followed by three washes in PBS-T, two washes in PBS, and one rinse in dH_2_O. Secondary antibodies were goat anti-rabbit Dylight 800 conjugate (1:2000, Thermo-Fisher, UK, catalogue #SA5-35571), goat anti-mouse Dylight 800 conjugate (1:2000, Thermo-Fisher, UK, catalogue #SA5-35521) and IRDye anti-mouse 800CW (1:2000, IRDye Licor Biosciences Ltd, UK, catalogue #926-32210). Membranes were imaged on an Odyssey Infrared Imager (LiCOR). The intensity of protein bands was quantified using Odyssey Image Studio software. Protein levels were determined and normalised to the level of the endogenous control.

### Immunofluorescence staining and confocal microscopy

Monolayer cultured cells were grown on coverslips until 70% confluent. Coverslips were washed three times with PBS and cells were fixed in PBS containing 5% formaldehyde at room temperature for 10 min. Permeabilization was carried out with PBS containing 0.5% NP40 at room temperature. Coverslips were then washed three times with PBS. Blocking was done in 10% filtered donkey serum in PBS for 30 min, followed by incubation with primary antibodies diluted in the blocking solution for 60 min at room temperature, and then washed three times with PBS followed by another wash in distilled H_2_O. Primary antibodies were SRSF1 (1:250, Mab96, Thermo Fisher Scientific, catalogue # 324500), SRSF2 (1:500, Abcam, catalogue #Ab204916), SRSF3 (1:300, Life Technologies, UK, catalogue #334200), HPV16 L1 (1:300 Dako, Denmark, clone K1H8), HPV16 E2 sheep antibody was a kind gift from Prof. Jo Parish, University of Birmingham, UK and was used at a dilution of 1:300, HPV E4 antibody clone B11 was a kind gift from Prof John Doorbar, University of Cambridge and was used at a dilution of 1:300. DAPI and secondary antibodies were diluted in blocking solution and added to the cells for 60 min, protected from light, prior to six washes in PBS followed by one wash in dH_2_O. Alexa fluor secondary anti-rabbit antibody 555 (1:1000, Invitrogen, UK, catalogue #A-31572), Alexa fluor secondary anti-mouse antibody 488 (1:1000, Invitrogen, UK, catalogue #A-21202), Alexa fluor secondary anti-sheep 488 antibody (1:1000, Invitrogen, UK, catalogue #A-11-15). Coverslips were mounted on glass slides with a glycerol-based mounting medium (Citifluor, AF1) and sealed with nail enamel. Samples were examined using a Zeiss LSM 710 confocal microscope and Zen black software (Zeiss) was used for capturing images.

Immunofluorescence staining of 3D organotypic raft culture sections was carried out on paraffin embedded fixed 2.5 µm sections. Antigen retrieval of the fixed section was performed by the histopathology department, Veterinary Diagnostic Services, University of Glasgow using heat induced epitope retrieval (HIER) using the Menarini Access Retrieval Unit method. Sample blocking, staining, and examination were conducted as described above.

### RNA extraction and cDNA synthesis

Total mRNA extraction was performed using TRIZOL reagent (Invitrogen, UK) according to the manufacturer’s instructions. For cells in monolayer culture, medium was removed, and cells were washed twice with PBS. Cells were lysed by scraping into 500 µl TRIZOL reagent for each well of a 6 well plate. Purified RNA was dissolved in DEPC-treated H_2_O and stored at −20°C. SuperScript^TM^ III First-Strand synthesis system for RT-PCR (Invitrogen, UK) was used for cDNA synthesis according to manufacturer’s instructions using (dT)_20_ primers.

### End point RT-PCR

For a single round of amplification 500 ng of cDNA was used in a total volume reaction of 20 µl using 200nM primers, 200 µM dNTPs, 1.5 mM MgCl_2_ and 2 units *Taq* polymerase (Invitrogen, UK). The amplification protocol was one cycle at 94° C for 2 min followed by 35 cycles of 94° C for 30 sec, annealing temperature of 55 ° C (E4^L1: E4 Forward primer 5’-GTTGTTGCACAGAGACTCAGTGG-3’ with L1 Reverse primer 5’-CGTGCAACATATTCATCCGTGC-3’) or annealing temperature of 57.8 ° C (E6E7: E6 Forward primer 5’-GAGAACTGCAATGTTTCAGGACCC-3’ with E7 Reverse primer 5’-GAACAGATGGGGCACACAATTCC-3’) for 45 sec followed by extension of 72° C for 1 min. There was a final extension cycle of 72° C for 10 min. GAPDH cDNA was amplified as an internal control (Forward primer 5’-AGGAAATGAGCTTGACAAAG-3’ and reverse primer 5’-ACCACAGTCCATGCCATCAC-3’)

### Nested RT-PCR

Amplification of L2L1 readthrough and E2 spliced mRNAs required two rounds of amplification. For L2L1, the first round of amplification used the same PCR reaction conditions described above with one cycle at 94° C for 2 min followed by 35 cycles of 94° C for 30 sec, annealing temperature of 57 ° C for 45 sec followed by an extension of 72° C for 1 min. There was a final extension cycle of 72° C for 10 min (L2 forward primer 5’-ACATGCAGCCTCACCTACTT-3’ with L1 reverse primer 1 5’-AACACCTACACAGGCCCAAA-3’). For the second round of amplification, 1µl of the first round PCR product was diluted in a total volume of 60 µl containing 200nM primers, 200 µM dNTPs, 1.5 mM MgCl_2_ and 2 units *Taq* polymerase. The PCR conditions were one cycle at 94°C for 2 min followed by 25 cycles of 94°C for 30 sec, annealing at 57°C for 45 sec and 72 °C extension for 1 min. This was followed with a final extension of 72 °C extension for 10 min (L2 forward primer 5’-ACATGCAGCCTCACCTACTT-3’ with L1 reverse primer 2 5’-TGTCCAACTGCAAGTAGTCTGGATGTTCCT-3’). For E2 cDNA the conditions for the first round of amplification were the same as for L2L1 cDNA except using E1 Forward primer 5’-ATCTACCATGGCTGATCCTGC-3’ and E2 Reverse primer 5’-GCAGTGAGGATTGGAGCACTGTC-3’. The conditions for the second round of amplification were the same as for L2L1 cDNA except that the annealing temperature was 63.2 °C for 35 cycles of amplification using E1^E2 cross splice Forward primer 5’-TGGCTGATCCTGCAGATTC-3’ and E2 Reverse primer 5’-CTTCGGTGCCCAAGGCGACGGCTTTGGTAT-3’. 10 µl of the final PCR products were fractionated on a 6% acrylamide gel. Electrophoresis was performed at 100 V and the gel was stained with 0.5 µg/ml EtBr for 15 min. PCR products were visualized under UV.

### Statistical analysis

Microsoft Excel software was used for plotting the data and calculating mean, standard deviations (SD), and p-values. Statistical significance was determined by students t-test with p≤L0.05 considered significant. All experiments were performed at least three times.

### Next Generation Sequencing

RNA was prepared using the Qiagen RNeasy kit from monolayer cultured NIKS16 cells grown for eight days to high density in the presence of 1.2. mM Ca^2+^ and treated either with DMSO or 10 µM SRPIN340 dissolved in DMSO for 48 hours. RNA quantification and quality was assessed by Qubit and Agilent TapeStation, respectively. Ribosomal RNA was depleted using the Illumina RiboZero Gold H/M/R Kit (Illumina, UK). Libraries were prepared using the TruSeq Stranded mRNA Kit (Illumina, UK) and cDNA was made using a Superscript II reverse transcriptase reaction (Invitrogen, UK). cDNA was converted to dsDNA and cleaned up using Beckman Coulter’s RNAClean XP (1.8 ratio) and then A-tailed. Illumina TruSeq LT adapters were ligated to the samples and cleaned up with a 1:1 bead ratio of AMPure XP followed by size selection using SPRIselect (Beckman Coulter, UK). The library was assessed for quality by Qubit and Agilent TapeStation. RNA paired-end sequencing was carried out using Illumina Genome Analyzer sequencing at the University of Glasgow.

### Alignment of fastq files and read summarization

The six raw RNA sequencing fastq files (3 x SRPIN340-treated and 3 x DMSO-treated) were analysed for quality using FastQC (version 0.11.2) and reads were trimmed of adaptor sequences and low-quality bases using Trimgalore. The trimmed reads were aligned to the human genome GRCh38 (*Ensembl*) using Hisat2 (version 2.0.5) [55] in the Linux environment (version 18.04.3). FeatureCounts (Version 1.5.1) [56] was used to quantify the reads by read summarization.

### Differential gene expression analysis

Differential gene expression analysis of SRPIN340-treated and DMSO-treated NIKS16 cells was determined using the Bioconductor package (Version 3.12) and DESeq2 (Version 1.30.1) [57], in the R environment (version 4.0.5). Differentially expressed genes were filtered by Q-value<0.05 and log fold change cut off of 2 except for comparison of SRPIN340-related versus HPV16 infection-related changes to NIKS16 gene expression where log fold change of 1.5 was used. Genes which were differentially expressed were variance stabilisation transformed for normalisation and visualisation on a heatmap. Pheatmap (version 1.0.12) was used for visualisation.

### Pathway Analysis

BiomaRt (version as 2.46.3) was used to convert *ensembl* gene ID into gene symbols before pathway analysis was performed. Over representation analysis was performed using ClusterProfiler (version 3.18.1) [58] in the R environment. Enrichment of gene ontology (GO) terms was determined for the subontology category, biological processes for 354 up and 241 down regulated genes (log fold change >2). The statistical test used includes a hypergeometric statistical test with Bonferroni correction. The thresholds used were Q-value < 0.05. Functional clustering of GO terms was performed using enrichplot (version 1.12.0).

### Comparison of differentially expressed genes upon HPV16 infection

Differentially expressed genes from HPV16-infected NIKS16 cells and uninfected NIKS cells [35] were compared manually to differentially expressed genes upon SRPIN340 treatment. Genes involved in epithelial barrier function and innate immunity were identified from both data sets.

### Over representation analysis of alternative splicing events

Bam files from the RNA paired-end sequencing of 10 µM SRPIN340-treated or DMSO-treated NIKS16 cells were sorted by co-ordinate, indexed and subject to SplAdder analysis [59] to measure and quantify alternative splicing events. Percentage spliced in (PSI) values were quantified for each splicing event and an independent two-tailed students t-test was performed for DMSO-treated and SRPIN340-treated values in order to determine the most significant differentially spliced genes upon SRPIN340 treatment. Pathway analysis was performed using Webgestalt ((http://www.webgestalt.org/) [39]. Over representation analysis was carried out testing for biological processes using Benjamini-Hochberg multiple testing adjustment.

## ACKNOWLEDGEMENTS

We would like to thank Prof. John Doorbar, University of Cambridge, UK, for providing NIKS and NIKS16 cells lines and the anti-E4 B11 antibody. The sheep anti-E2 antibody was a kind gift from Prof. Joanna Parish, University of Birmingham, UK. Our thanks go to Dr Jean Quinn, University of Glasgow, for 3D raft culture training.

## DATA AVAILABILITY STATEMENT

The raw data for the results described here is available upon request. RNA-Seq data sets are freely available from the European Nucleotide Archive accession number PRJEB80489.

## AUTHOR CONTRIBUTIONS

AAA. Faizo: Investigation, formal analysis, validation and visualization, writing review and editing.

C. Bellward: Investigation, formal analysis, validation and visualization.

H. Hernandez Lopez: Investigation, formal analysis, validation and visualization, writing review and editing.

A. Stevenson: methodology, supervision, project administration.

Gu Q.: Investigation, formal analysis, supervision, writing review and editing.

S. Graham: Conceptualization, supervision, writing, review and editing

---

**Supplementary Figure 1. NIKS16 cells contain episomal genomes and undergo late events in the viral life cycle.** A. Southern blot analysis of HPV16 genomes in NIKS16 cells. DNA was isolated from monolayer cultured NIKS16 cells and digested with either Bam HI, which cuts the genome once (lane 2), or Hind III, which does not cut the genome (lane 3). The HPV16 genome was cut from plasmid pEF-HPV16 with Bam HI as a size control for the episomal genome (lane 1). B. Western blot analysis of HPV16 E2 and L1 expression in protein lysates from NIKS and NIKS16 cells grown in monolayer culture to give an undifferentiated cell population (U, lanes 1 & 2) or differentiated by growing to high density in the presence of 1.2 mM Ca^2+^ (D, lanes 3 & 4). Involucrin and keratin 10 were used as controls for keratinocyte differentiation. GADPH is used as a loading control.

**Supplementary Figure 2. Topical SRPIN340 treatment of 3D raft tissues does not disrupt tissue morphology and may be transmitted throughout the tissues.** A. H&E staining of NIKS16 3D raft tissues grown for 12 days then grown in the presence of 10 or 50 µM SRPIN340 in the culture medium for 48 hours. B. H&E staining of NIKS16 3D raft tissues grown for 12 days followed by topical application of 10 or 50 µM SRPIN340 in a 50 µl droplet and incubation for 48 hours. C. Autofluorescence control of NIKS16 3D tissues stained with the same secondary antibody used in D. D. Immunofluorescence microscopy of NIKS16 3D tissues topically treated with Alexa-fluor 488 (Alexa488: green staining). Tissues in the right-hand panels of C. and D. are counterstained with DAPI. The junction of the basal layer of cells with the dermal equivalent is indicated by a dotted line. Size bars = 50 µM.

**Supplementary Figure 3. SRPIN340 treatment does not alter expression of the HPV16 major late mRNAs.** A. Diagram of the portion of the HPV16 genome downstream of the P_670_ promoter. The E1^E4^L1 alternatively spliced mRNA that encodes E4 and L1 and the readthrough mRNA E1^E4, E5, L2 and L1 that is thought to encode L2 are shown beloiw the genome map. P_670_, HPV16 late promoter located at nucleotide 670. p(A)L, late polyadenylation site. Open rectangles, HPV16 genes E1, E2, E4, E5, L2 and L1. Shaded rectangles, open reading frames. Lines represent RNA sequences which are spliced out to form the mRNAs. AAAn, poly(A) tail. Chevrons indicate the approximate positions of forward and reverse primers used in the PCR reactions shown in (B) and (C). B. Ethidium bromide-stained gel showing the PCR products obtained from amplification of NIKS16 cDNA using the primer pair indicated in (A) for the E1^E4^L1 mRNA. GAPDH primer pairs were added to the same reaction for a loading control. * non-specific bands. C. Upper panel: ethidium bromide-stained gel showing the PCR products obtained from amplification of NIKS16 cDNA using the primer pairs indicated in (A) in a nested PCR reaction to amplify across the L2-L1 junction from the E1^E4, E5, L2, L1 readthrough mRNA. Lower panel: amplification of GAPDH as a loading control. In (B) and (C) the gel pictures are split because additional reactions were electrophoresed between the marker lane and the DMSO and SRPIN340 lanes.

## References

1. Egawa N, Egawa K, Griffin H, Doorbar J. Human Papillomaviruses; Epithelial Tropisms, and the Development of Neoplasia. Viruses. 2015;7(7):2802. PubMed PMID: doi:10.3390/v7072802.

2. Stanley MA. Epithelial Cell Responses to Infection with Human Papillomavirus. Clin Microbiol Rev. 2012;25(2):215–22. doi: 10.1128/cmr.05028-11.

3. Graham SV. The human papillomavirus replication cycle, and its links to cancer progression: a comprehensive review. Clin Sci (Lond). 2017;131(17):2201–21. Epub 20170810. doi: 10.1042/cs20160786.

4. Harper DM, DeMars LR. HPV vaccines – A review of the first decade. Gynecol Oncol. 2017;146(1):196–204. doi: 10.1016/j.ygyno.2017.04.004.

5. Kirk A, Graham SV. The human papillomavirus late life cycle and links to keratinocyte differentiation. J Med Virol. 2024;96(2):e29461. doi: 10.1002/jmv.29461.

6. McBride AA. Human papillomaviruses: diversity, infection and host interactions. Nat Rev Microbiol. 2022;20(2):95–108. Epub 20210914. doi: 10.1038/s41579-021-00617-5.

7. Vande Pol SB, Klingelhutz AJ. Papillomavirus E6 oncoproteins. Virology. 2013;445(1–2):115–37. doi: 10.1016/j.virol.2013.04.026.

8. Roman A, Munger K. The papillomavirus E7 proteins. Virology. 2013;445(1):138–68. doi: 10.1016/j.virol.2013.04.013.

9. Egawa N, Wang Q, Griffin HM, Murakami I, Jackson D, Mahmood R, Doorbar J. HPV16 and 18 genome amplification show different E4-dependence, with 16E4 enhancing E1 nuclear accumulation and replicative efficiency via its cell cycle arrest and kinase activation functions. PLOS Pathogens. 2017;13(3):e1006282. doi: 10.1371/journal.ppat.1006282.

10. Zhou Z, Fu X-D. Regulation of splicing by SR proteins and SR protein-specific kinases. Chromosoma. 2013;122(3):191–207. doi: 10.1007/s00412-013-0407-z.

11. Long JC, Caceres JF. The SR protein family of splicing factors: master regulators of gene expression. Biochem J. 2009;417(1):15–27.

12. Giannakouros T, Nikolakaki E, Mylonis I, Georgatsou E. Serine-arginine protein kinases: a small protein kinase family with a large cellular presence. FEBS J. 2011;278(4):570–86. doi: 10.1111/j.1742-4658.2010.07987.x.

13. Aubol BE, Wu G, Keshwani MM, Movassat M, Fattet L, Hertel KJ, et al. Release of SR Proteins from CLK1 by SRPK1: A Symbiotic Kinase System for Phosphorylation Control of Pre-mRNA Splicing. Mol Cell. 2016;63(2):218–28. Epub 20160707. doi: 10.1016/j.molcel.2016.05.034.

14. Yun CY, Fu XD. Conserved SR protein kinase functions in nuclear import and its action is counteracted by arginine methylation in Saccharomyces cerevisiae. J Cell Biol. 2000;150(4):707–18. doi: 10.1083/jcb.150.4.707.

15. Lai MC, Lin RI, Huang SY, Tsai CW, Tarn WY. A human importin-beta family protein, transportin-SR2, interacts with the phosphorylated RS domain of SR proteins. J Biol Chem. 2000;275(11):7950–7. doi: 10.1074/jbc.275.11.7950.

16. Nikolakaki E, Sigala I, Giannakouros T. Good Cop, Bad Cop: The Different Roles of SRPKs. Front Genet. 2022;13:902718. Epub 20220602. doi: 10.3389/fgene.2022.902718.

17. Graham SV. Human papillomavirus: gene expression, regulation and prospects for novel diagnostic methods and antiviral therapies. Future Microbiol. 2010;5(10):1493–505.

18. Yu L, Majerciak V, Zheng ZM. HPV16 and HPV18 Genome Structure, Expression, and Post-Transcriptional Regulation. Int J Mol Sci. 2022;23(9). Epub 20220429. doi: 10.3390/ijms23094943.

19. Klymenko T, Hernandez-Lopez H, MacDonald AI, Bodily JM, Graham SV. Human papillomavirus E2 regulates SRSF3 (SRp20) to promote capsid protein expression in infected differentiated keratinocytes. J Virol. 2016. doi: 10.1128/jvi.03073-15.

20. Mole S, Faizo AAA, Hernandez-Lopez H, Griffiths M, Stevenson A, Roberts S, Graham SV. Human papillomavirus type 16 infection activates the host serine arginine protein kinase 1 (SRPK1) - splicing factor axis. J Gen Virol. 2020;101(5):523–32. Epub 20200313. doi: 10.1099/jgv.0.001402.

21. Bodaghi S, Jia R, Zheng Z-M. Human papillomavirus type 16 E2 and E6 are RNA-binding proteins and inhibit in vitro splicing of pre-mRNAs with suboptimal splice sites. Virology. 2009;386(1):32–43. doi: 10.1016/j.virol.2008.12.037.

22. Oltean S, Gammons M, Hulse R, Hamdollah-Zadeh M, Mavrou A, Donaldson L, et al. SRPK1 inhibition in vivo: modulation of VEGF splicing and potential treatment for multiple diseases. Biochem Soc Trans. 2012;40(4):831–5. doi: 10.1042/bst20120051.

23. Bates DO, Morris JC, Oltean S, Donaldson LF. Pharmacology of Modulators of Alternative Splicing. Pharmacol Rev. 2017;69(1):63–79. doi: 10.1124/pr.115.011239.

24. Karakama Y, Sakamoto N, Itsui Y, Nakagawa M, Tasaka-Fujita M, Nishimura-Sakurai Y, et al. Inhibition of hepatitis C virus replication by a specific inhibitor of serine-arginine-rich protein kinase. Antimicrob Agents Chemother. 2010;54(8):3179–86. Epub 20100524. doi: 10.1128/aac.00113-10.

25. Fukuhara T, Hosoya T, Shimizu S, Sumi K, Oshiro T, Yoshinaka Y, et al. Utilization of host SR protein kinases and RNA-splicing machinery during viral replication. Proc Natl Acad Sci U S A. 2006;103(30):11329–33. Epub 20060713. doi: 10.1073/pnas.0604616103.

26. Sciabica KS, Dai QJ, Sandri-Goldin RM. ICP27 interacts with SRPK1 to mediate HSV splicing inhibition by altering SR protein phosphorylation. EMBO J. 2003;22(7):1608–19.

27. Daub H, Blencke S, Habenberger P, Kurtenbach A, Dennenmoser J, Wissing J, et al. Identification of SRPK1 and SRPK2 as the major cellular protein kinases phosphorylating hepatitis B virus core protein. J Virol. 2002;76(16):8124–37. doi: 10.1128/jvi.76.16.8124-8137.2002.

28. Kajitani N, Schwartz S. The role of RNA-binding proteins in the processing of mRNAs produced by carcinogenic papillomaviruses. Semin Cancer Biol. 2022;86(Pt 3):482–96. Epub 20220216. doi: 10.1016/j.semcancer.2022.02.014.

29. Graham SV. HPV and RNA Binding Proteins: What We Know and What Remains to Be Discovered. Viruses. 2024;16(5). Epub 20240515. doi: 10.3390/v16050783.

30. Weschler EI, Wang Q, Roberts I, Pagliarulo E, Jackson D, Untersperger C, et al. Reconstruction of human papillomavirus type 16-mediated early-stage neoplasia implicated E6/E7 deeregualtion and the loss of contact inhibition in neoplastic progression. J Virol. 2012;86(11):6358–64.

31. Flores ER, Allen-Hoffmann BL, Lee D, Sattler CA, Lambert PF. Establishment of the human papillomavirus type 16 (HPV-16) life cycle in an immortalized human foreskin keratinocyte cell line. Virology. 1999;262. doi: 10.1006/viro.1999.9868.

32. Weber PA, Chang HC, Spaeth KE, Nitsche JM, Nicholson BJ. The permeability of gap junction channels to probes of different size is dependent on connexin composition and permeant-pore affinities. Biophys J. 2004;87(2):958–73. doi: 10.1529/biophysj.103.036350.

33. Zheng Y, Cui X, Nilsson K, Yu H, Gong L, Wu C, Schwartz S. Efficient production of HPV16 E2 protein from HPV16 late mRNAs spliced from SD880 to SA2709. Virus Res. 2020;285:198004. Epub 20200505. doi: 10.1016/j.virusres.2020.198004.

34. Evans MR, James CD, Bristol ML, Nulton TJ, Wang X, Kaur N, et al. Human Papillomavirus 16 E2 Regulates Keratinocyte Gene Expression Relevant to Cancer and the Viral Life Cycle. J Virol 2019;93(4):e01941–18. doi: 10.1128/JVI.01941-18 %J.

35. Klymenko T, Gu Q, Herbert I, Stevenson A, Iliev V, Watkins G, et al. RNA-Seq Analysis of Differentiated Keratinocytes Reveals a Massive Response to Late Events during Human Papillomavirus 16 Infection, Including Loss of Epithelial Barrier Function. J Virol 2017;91(24):e01001–17. doi: 10.1128/JVI.01001-17 %J.

36. Matsui T, Amagai M. Dissecting the formation, structure and barrier function of the stratum corneum. Int Immunol. 2015;27(6):269–80. doi: 10.1093/intimm/dxv013.

37. Lessard JC, Piña-Paz S, Rotty JD, Hickerson RP, Kaspar RL, Balmain A, Coulombe PA. Keratin 16 regulates innate immunity in response to epidermal barrier breach. Proc Natl Acad Sci U S A. 2013;110(48):19537–42. Epub 20131111. doi: 10.1073/pnas.1309576110.

38. Carregaro F, Stefanini ACB, Henrique T, Tajara EH. Study of small proline-rich proteins (SPRRs) in health and disease: a review of the literature. Arch Dermatol Res. 2013;305(10):857–66. doi: 10.1007/s00403-013-1415-9.

39. Liao Y, Wang J, Jaehnig EJ, Shi Z, Zhang B. WebGestalt 2019: gene set analysis toolkit with revamped UIs and APIs. Nucleic Acids Res. 2019;47(W1):W199-w205. doi: 10.1093/nar/gkz401.

40. Cartegni L, Wang J, Zhu Z, Zhang MQ, Krainer AR. ESEfinder: A web resource to identify exonic splicing enhancers. Nucleic Acids Res. 2003;31(13):3568–71. doi: 10.1093/nar/gkg616.

41. Hao C, Zheng Y, Jönsson J, Cui X, Yu H, Wu C, et al. hnRNP G/RBMX enhances HPV16 E2 mRNA splicing through a novel splicing enhancer and inhibits production of spliced E7 oncogene mRNAs. Nucleic Acids Res. 2022;50(7):3867–91. doi: 10.1093/nar/gkac213.

42. Gauson EJ, Windle B, Donaldson MM, Caffarel MM, Dornan ES, Coleman N, et al. Regulation of human genome expression and RNA splicing by human papillomavirus 16 E2 protein. Virology. 2014;468–470:10-8. doi: 10.1016/j.virol.2014.07.022.

43. Prescott EL, Brimacombe CL, Hartley M, Bell I, Graham S, Roberts S. Human Papillomavirus Type 1 E1E4 Protein Is a Potent Inhibitor of the Serine-Arginine (SR) Protein Kinase SRPK1 and Inhibits Phosphorylation of Host SR Proteins and of the Viral Transcription and Replication Regulator E2. J Virol. 2014;88(21):12599–611. doi: 10.1128/jvi.02029-14.

44. Patel M, Sachidanandan M, Adnan M. Serine arginine protein kinase 1 (SRPK1): a moonlighting protein with theranostic ability in cancer prevention. Mol Biol Rep. 2019;46(1):1487–97. Epub 20181208. doi: 10.1007/s11033-018-4545-5.

45. Johansson C, Somberg M, Li X, Backström Winquist E, Fay J, Ryan F, et al. HPV-16 E2 contributes to induction of HPV-16 late gene expression by inhibiting early polyadenylation. EMBO J. 2012;31(14):3212–27. Epub 20120522. doi: 10.1038/emboj.2012.147.

46. Davy C, McIntosh P, Jackson DJ, Sorathia R, Miell M, Wang Q, et al. A novel interaction between the human papillomavirus type 16 E2 and E1E4 proteins leads to stabilization of E2. Virology. 2009;394(2):266–75. doi: 10.1016/j.virol.2009.08.035.

47. Siddiqa A, Léon KC, James CD, Bhatti MF, Roberts S, Parish JL. The human papillomavirus type 16 L1 protein directly interacts with E2 and enhances E2-dependent replication and transcription activation. J Gen Virol. 2015;96(8):2274–85. doi: doi:10.1099/vir.0.000162.

48. Bazzi H, Christiano AM. Broken hearts, woolly hair, and tattered skin: when desmosomal adhesion goes awry. Curr Opin Cell Biol. 2007;19(5):515–20. doi:10.1016/j.ceb.2007.08.001.

49. Morizane S, Sunagawa K, Nomura H, Ouchida M. Aberrant serine protease activities in atopic dermatitis. J Dermatol Sci. 2022;107(1):2–7. Epub 20220628. doi: 10.1016/j.jdermsci.2022.06.004.

50. Kanno Y, Matsui Y. Cellular coupling in cancerous stomach epithelium. Nature. 1968;218:775–6.

51. Guo T, Zambo KDA, Zamuner FT, Ou T, Hopkins C, Kelley DZ, et al. Chromatin structure regulates cancer-specific alternative splicing events in primary HPV-related oropharyngeal squamous cell carcinoma. Epigenetics. 2020;15(9):959–71. Epub 20200322. doi: 10.1080/15592294.2020.1741757.

52. Kelley DZ, Flam EL, Guo T, Danilova LV, Zamuner FT, Bohrson C, et al. Functional characterization of alternatively spliced GSN in head and neck squamous cell carcinoma. Transl Res. 2018;202:109–19. Epub 20180726. doi: 10.1016/j.trsl.2018.07.007.

53. Jeon S, Allen-Hoffman BL, Lambert PF. Integration of human papillomavirus type 16 into the human genome correlates with a selective growth advantage of cells. J Virol. 1995;69(5):2989–97.

54. Anacker D, Moody C. Generation of organotypic raft cultures from primary human keratinocytes. J Vis Exp. 2012;(60). Epub 20120222. doi: 10.3791/3668.

55. Kim D, Paggi JM, Park C, Bennett C, Salzberg SL. Graph-based genome alignment and genotyping with HISAT2 and HISAT-genotype. Nat Biotechnol. 2019;37(8):907–15. Epub 20190802. doi: 10.1038/s41587-019-0201-4.

56. Liao Y, Smyth GK, Shi W. featureCounts: an efficient general purpose program for assigning sequence reads to genomic features. Bioinformatics. 2014;30(7):923–30. Epub 20131113. doi: 10.1093/bioinformatics/btt656.

57. Love MI, Huber W, Anders S. Moderated estimation of fold change and dispersion for RNA-seq data with DESeq2. Genome Biol. 2014;15(12):550. doi: 10.1186/s13059-014-0550-8.

58. Yu G, Wang LG, Han Y, He QY. clusterProfiler: an R package for comparing biological themes among gene clusters. Omics. 2012;16(5):284–7. Epub 20120328. doi: 10.1089/omi.2011.0118.

59. Kahles A, Ong CS, Zhong Y, Rätsch G. SplAdder: identification, quantification and testing of alternative splicing events from RNA-Seq data. Bioinformatics. 2016;32(12):1840–7. Epub 20160211. doi: 10.1093/bioinformatics/btw076.

