## Supplementary figures and images for "The splicing factor kinase, SR protein kinase 1 (SRPK1) is essential for late events in the human papillomavirus life cycle"

### Supplemental Figure 1

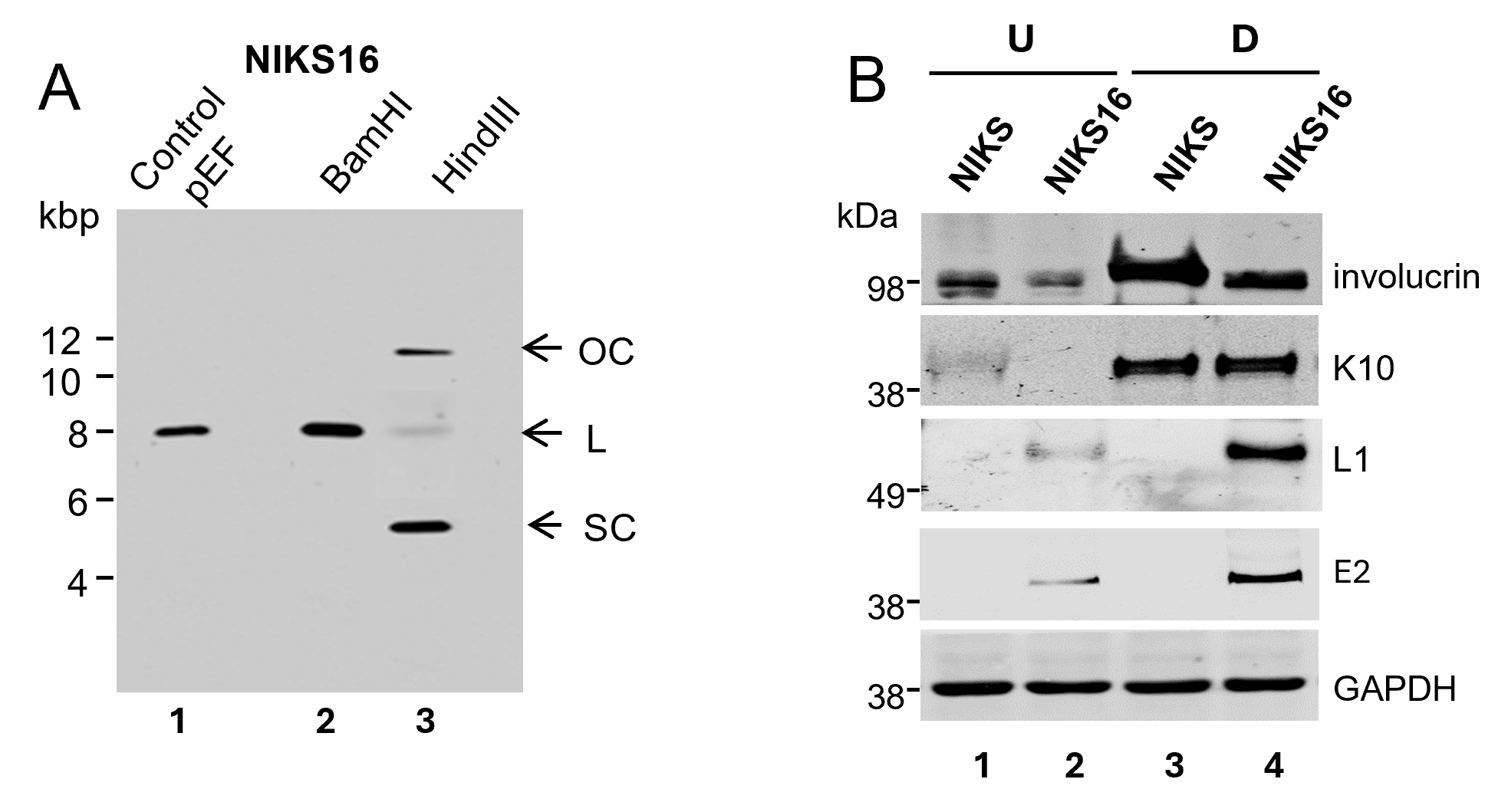

### Supplemental Figure 2

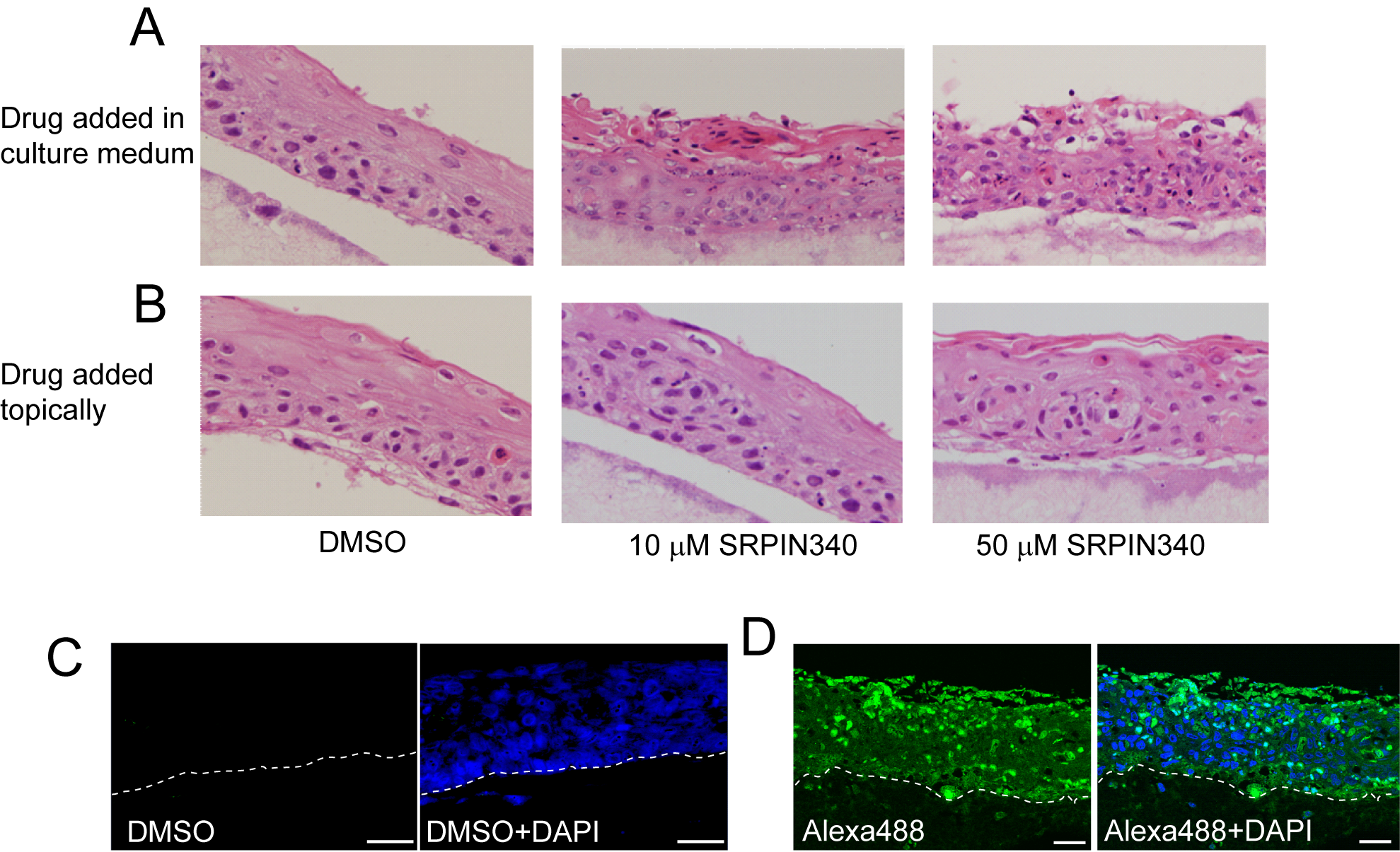

### Supplemental Figure 3

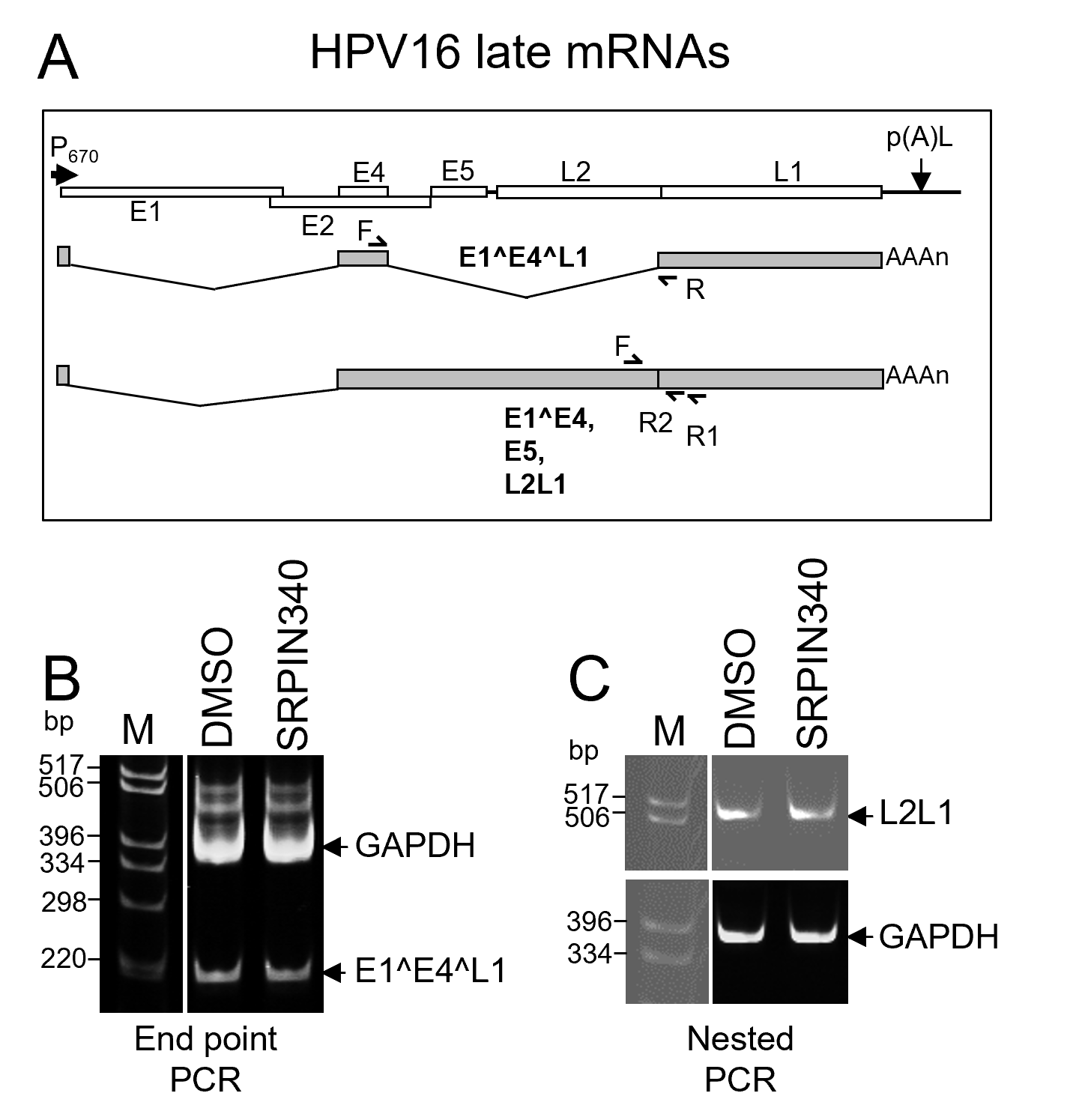
