## Supplemental Table 1 for "The splicing factor kinase, SR protein kinase 1 (SRPK1) is essential for late events in the human papillomavirus life cycle"

**Supplementary Table 1.** List of genes differentially expressed upon SRPIN340 treatment determined by DESeq2. Q-value <0.05.

| Gene | log2FoldChange | Q-value |
| --- | --- | --- |
| TMPRSS11D | 6.797647766 | 8.66E-05 |
| SERPINB11 | 6.032431007 | 0.000679 |
| CYP1A1 | 5.968312781 | 1.21E-25 |
| HRNR | 5.415118392 | 0.004903 |
| ENDOU | 4.90915265 | 1.9E-06 |
| LOR | 4.607461151 | 0.001384 |
| FAIM2 | 4.555164059 | 0.010715 |
| LIPN | 4.437793821 | 0.017848 |
| SH2D1B | 4.393057631 | 2.56E-05 |
| CRNN | 4.163890496 | 6.37E-29 |
| CAPN14 | 4.084787111 | 1.06E-05 |
| AQP5 | 4.069593091 | 6.13E-19 |
| KLK12 | 4.00172104 | 4.12E-13 |
| TMPRSS11A | 3.98695018 | 3.71E-15 |
| FLG2 | 3.887629517 | 9.25E-17 |
| CEACAM5 | 3.819780979 | 0.000152 |
| AC025154.2 | 3.680746053 | 1.83E-13 |
| S100A7 | 3.52549549 | 1.83E-50 |
| ARG1 | 3.473623525 | 0.000287 |
| SPINK7 | 3.463702469 | 1.75E-31 |
| AC005747.1 | 3.419194555 | 0.021177 |
| SPAG17 | 3.4159458 | 8.97E-05 |
| SPRR3 | 3.345138016 | 5.65E-20 |
| S100A12 | 3.278182015 | 2.07E-14 |
| ACER1 | 3.228005898 | 0.038789 |
| CYP1B1 | 3.173612714 | 2.76E-12 |
| SPINK5 | 3.118849357 | 4.06E-35 |
| KRTDAP | 3.110464939 | 1.35E-14 |
| CEACAM6 | 3.081712305 | 2.4E-14 |
| FAM83C | 3.081258642 | 5.77E-05 |
| FLG | 3.074589309 | 3.43E-43 |
| PLA2G4D | 3.032090612 | 0.006545 |
| SNX31 | 2.994435232 | 0.046948 |
| CHI3L1 | 2.985669305 | 0.015534 |
| GPR78 | 2.967342661 | 0.000542 |
| RDH12 | 2.964841315 | 2.75E-10 |
| FABP4 | 2.958742078 | 0.000823 |
| LIPM | 2.946469386 | 0.023147 |
| WFDC12 | 2.944169304 | 0.000446 |
| CALB1 | 2.861319402 | 8.97E-05 |
| FGG | 2.83556582 | 0.017908 |
| LINC00589 | 2.828017185 | 2.44E-09 |
| RNF224 | 2.820035347 | 0.027623 |
| S100A8 | 2.796318445 | 6.91E-18 |
| PADI1 | 2.75167383 | 0.035897 |
| S100A9 | 2.716664496 | 3.01E-10 |
| NMRAL2P | 2.687185356 | 0.009493 |
| HMGCS2 | 2.672135661 | 0.030678 |
| HAL | 2.616531474 | 2.75E-10 |
| CYP4F12 | 2.608421058 | 0.000261 |
| DSG1 | 2.598191597 | 1.34E-25 |
| BMP7 | 2.588187415 | 0.000366 |
| MAL | 2.580201362 | 0.000287 |
| SERPINB4 | 2.542729569 | 2.3E-18 |
| CYP2C18 | 2.506529455 | 0.046326 |
| SERPINB3 | 2.48433339 | 5.95E-16 |
| CDH16 | 2.478314162 | 9.89E-17 |
| GPLD1 | 2.429878197 | 6.36E-07 |
| P2RX6P | 2.399665974 | 0.030153 |
| KYNU | 2.394760257 | 7E-08 |
| SLC40A1 | 2.350342495 | 0.001118 |
| RNF222 | 2.349196076 | 0.018527 |
| UPK1A | 2.347305669 | 0.012691 |
| ST6GALNAC1 | 2.340037292 | 1.6E-07 |
| TTC39A | 2.33687487 | 4.01E-05 |
| ACOX2 | 2.327367987 | 0.006724 |
| KRT78 | 2.323194682 | 1.12E-07 |
| KRT2 | 2.302088478 | 0.000736 |
| MYRF | 2.27371081 | 0.010069 |
| BCAS1 | 2.265109315 | 0.002092 |
| MUC15 | 2.249972714 | 2.1E-06 |
| SPTSSB | 2.242123871 | 2.88E-08 |
| PLA2G4E | 2.239230679 | 0.018743 |
| SERPINB12 | 2.236831591 | 9.07E-06 |
| CWH43 | 2.229531348 | 4.12E-12 |
| ASPG | 2.218561959 | 0.001697 |
| SYTL5 | 2.196324339 | 0.000643 |
| SDR16C5 | 2.1919528 | 1.66E-20 |
| PTPN22 | 2.152200098 | 0.001249 |
| IL36RN | 2.138558314 | 2.41E-05 |
| KRT77 | 2.123362668 | 2.08E-15 |
| CYP4F3 | 2.108318739 | 2.28E-07 |
| PTPRH | 2.107162611 | 0.016335 |
| CARD18 | 2.107122408 | 5.69E-19 |
| FAM3D | 2.104486001 | 0.005246 |
| GCNT3 | 2.099692087 | 1.42E-05 |
| C8orf89 | 2.084568716 | 0.046326 |
| SPRR1A | 2.069598699 | 3.69E-05 |
| CD86 | 2.06554242 | 2.65E-06 |
| SBSN | 2.048318457 | 1.43E-25 |
| DAPL1 | 2.038744052 | 3.79E-06 |
| MEOX1 | 2.025739639 | 0.030568 |
| XKRX | 2.023548742 | 0.00275 |
| DEGS2 | 2.011137052 | 0.026037 |
| HPGD | 2.004686543 | 3.23E-07 |
| VWF | 2.001526731 | 0.000684 |
| CPM | 2.001010661 | 0.002575 |
| ATP10B | 1.992530562 | 0.001875 |
| SLURP1 | 1.973454044 | 0.000287 |
| TEX19 | 1.971111547 | 0.001986 |
| KRT4 | 1.946330232 | 1.55E-05 |
| SULT2B1 | 1.917757555 | 0.000001 |
| BNIPL | 1.914493475 | 2.28E-10 |
| SCAT8 | 1.911406612 | 0.000146 |
| SLC6A14 | 1.909728824 | 7.91E-08 |
| SLC16A7 | 1.898371696 | 3.82E-09 |
| TMPRSS11F | 1.894430338 | 1.91E-06 |
| PI3 | 1.880215251 | 1.38E-09 |
| KRT1 | 1.872868602 | 2.71E-18 |
| EREG | 1.863569422 | 1.32E-10 |
| ADH7 | 1.862807067 | 0.001424 |
| ITGB7 | 1.846481665 | 2.17E-05 |
| ALDH3A1 | 1.812401173 | 0.001953 |
| IL1A | 1.806840171 | 2.35E-17 |
| MAB21L4 | 1.805892773 | 0.002527 |
| S100P | 1.802888864 | 0.000191 |
| C5orf46 | 1.799694563 | 2.17E-07 |
| AGR2 | 1.79507323 | 7.25E-05 |
| AC023906.5 | 1.794867855 | 0.046198 |
| SUSD4 | 1.794437401 | 0.000182 |
| PLA2G4F | 1.782133645 | 3.99E-09 |
| PBX1 | 1.762784796 | 5.32E-05 |
| KHDC1L | 1.762444083 | 0.047124 |
| PGLYRP3 | 1.755523903 | 0.000126 |
| ECM2 | 1.752211056 | 0.001908 |
| SPRR1B | 1.744230684 | 0.001421 |
| SLC39A2 | 1.716132516 | 0.015088 |
| PIR | 1.70161939 | 1.16E-14 |
| PGLYRP4 | 1.699943496 | 0.000193 |
| TMEM45A | 1.694195559 | 3.89E-16 |
| MPP7 | 1.690522756 | 6.08E-12 |
| IL1B | 1.689738451 | 5.15E-05 |
| GDA | 1.68822189 | 0.001472 |
| KRT10 | 1.686715106 | 1.91E-09 |
| WFDC21P | 1.667788812 | 0.003202 |
| ANKRD35 | 1.666211028 | 1.86E-07 |
| CSTA | 1.662282698 | 3.04E-07 |
| GALNT12 | 1.647562817 | 1.08E-05 |
| AL118508.1 | 1.636577952 | 0.000453 |
| LINC01559 | 1.631522728 | 5.91E-05 |
| MSMB | 1.627686802 | 0.026423 |
| FABP5P7 | 1.626466101 | 6.53E-11 |
| FBP1 | 1.608973526 | 0.020955 |
| VSIG10L | 1.608870209 | 1.65E-06 |
| ADGRF1 | 1.607114895 | 0.003991 |
| CD36 | 1.602547495 | 0.000287 |
| LCN2 | 1.595159587 | 0.015484 |
| MX2 | 1.587322776 | 0.011657 |
| NOD2 | 1.584115802 | 0.004549 |
| LINC00330 | 1.58252474 | 0.004059 |
| DSC1 | 1.580088623 | 4.08E-07 |
| N4BP3 | 1.575754909 | 0.029008 |
| LIPK | 1.572973154 | 0.000647 |
| ANKRD22 | 1.560027802 | 2.01E-07 |
| SLC27A2 | 1.559638977 | 8.54E-06 |
| CMAHP | 1.557481394 | 0.005941 |
| RPTN | 1.531215494 | 0.000571 |
| CCNA1 | 1.522164896 | 4.33E-14 |
| TMPRSS4 | 1.509295842 | 6.01E-06 |
| FABP5 | 1.499582531 | 1.27E-20 |
| ALDH3B2 | 1.496885954 | 0.000943 |
| CLCA4 | 1.486846305 | 0.00083 |
| PPM1K | 1.484238353 | 6.89E-07 |
| RAET1L | 1.473925662 | 0.000047 |
| ALOX15B | 1.470963277 | 1.96E-06 |
| GSTA4 | 1.470659363 | 2.36E-05 |
| ATP6V0A4 | 1.470597931 | 0.000601 |
| TM7SF2 | 1.470478994 | 1.08E-05 |
| SERPINB13 | 1.469905877 | 4.39E-12 |
| CA2 | 1.466043918 | 0.000165 |
| ADGRL2 | 1.457218457 | 9.71E-09 |
| ALDH5A1 | 1.447200484 | 5.64E-05 |
| CHAC1 | 1.446583538 | 0.003195 |
| GBP6 | 1.436590984 | 4.37E-22 |
| MYH14 | 1.433387497 | 0.020913 |
| NCF2 | 1.432012138 | 0.001592 |
| NDRG2 | 1.420031337 | 0.001494 |
| AP002954.2 | 1.417655891 | 0.021279 |
| LINC02178 | 1.40953805 | 0.048026 |
| CYP3A5 | 1.404366286 | 0.002257 |
| DHRS9 | 1.386657162 | 0.002576 |
| KRT79 | 1.386439432 | 0.042316 |
| IGFBP4 | 1.385664928 | 0.04145 |
| FLVCR2 | 1.377515881 | 0.001332 |
| SLC7A11 | 1.374361259 | 2.73E-07 |
| GLTP | 1.362957023 | 1.19E-10 |
| MAP3K8 | 1.351789707 | 0.044839 |
| AQP3 | 1.331027844 | 0.000643 |
| POF1B | 1.329122273 | 1.64E-06 |
| AC011374.1 | 1.321050784 | 0.003713 |
| POLR2J3 | 1.320014004 | 0.003869 |
| HMOX1 | 1.309826998 | 0.009557 |
| STX19 | 1.305352329 | 0.034124 |
| BLNK | 1.294668966 | 0.000943 |
| FUCA1 | 1.292616757 | 0.000678 |
| NMNAT3 | 1.285826226 | 0.021162 |
| DEFB1 | 1.281333917 | 0.004293 |
| KRT16 | 1.280428101 | 1.46E-07 |
| TTC9 | 1.273281111 | 0.002707 |
| BCO2 | 1.269185502 | 1.91E-06 |
| PKIB | 1.26434816 | 0.023019 |
| RASSF5 | 1.258600754 | 0.001424 |
| CAPNS2 | 1.258275705 | 0.00324 |
| ABCC2 | 1.258206966 | 0.001209 |
| SYTL4 | 1.254387173 | 0.045851 |
| GRAMD1C | 1.252536852 | 0.01101 |
| SIPA1L2 | 1.248603725 | 1.87E-07 |
| HBEGF | 1.247319854 | 0.000135 |
| USH1G | 1.247235112 | 0.003261 |
| SERPINB7 | 1.240907151 | 1.79E-05 |
| SLC44A5 | 1.232172353 | 4.96E-05 |
| FOXA1 | 1.229620242 | 0.014375 |
| RAB27B | 1.212624408 | 6.64E-05 |
| AL035661.1 | 1.20888795 | 0.035595 |
| KRT6B | 1.208606554 | 0.000278 |
| ZNF736P9Y | 1.185139201 | 0.000281 |
| WNT2B | 1.183917746 | 0.01206 |
| MYZAP | 1.183345883 | 0.016853 |
| ALDH1A3 | 1.171792679 | 0.000684 |
| EXOC6 | 1.170118128 | 0.006411 |
| GPX2 | 1.158765954 | 1.09E-06 |
| SCEL | 1.156697489 | 1.05E-08 |
| IGFL2 | 1.145213336 | 2.03E-07 |
| OAS1 | 1.14235205 | 0.007654 |
| MCTP1 | 1.132820779 | 0.019321 |
| GRHL1 | 1.131096638 | 1.15E-05 |
| TMPRSS13 | 1.130859648 | 0.044468 |
| LPAR6 | 1.124676384 | 0.001997 |
| CD164L2 | 1.122706245 | 0.021023 |
| KLK11 | 1.117067353 | 0.000379 |
| KAZALD1 | 1.115794854 | 0.023019 |
| RNF128 | 1.115585357 | 2.62E-07 |
| ADTRP | 1.106172042 | 0.000158 |
| CCDC18-AS1 | 1.104416582 | 0.000104 |
| RAET1E | 1.093243575 | 8.13E-05 |
| GCOM1 | 1.084070974 | 0.000774 |
| TGM1 | 1.07432035 | 0.016362 |
| ABCA12 | 1.04633919 | 1.16E-10 |
| P2RY1 | 1.039832806 | 0.001311 |
| HINT3 | 1.037578204 | 0.001895 |
| MBOAT2 | 1.03457057 | 1.91E-06 |
| ACPP | 1.034174388 | 0.025619 |
| CTH | 1.031777268 | 0.045492 |
| CREG2 | 1.020224094 | 0.016053 |
| NGFR | 1.007210347 | 1.54E-05 |
| ANKRD1 | -4.307959272 | 0.000146 |
| MYL9 | -4.205082349 | 0.000063 |
| MYL7 | -3.538350233 | 2.91E-06 |
| GLP2R | -3.287298644 | 0.021162 |
| LINC00052 | -2.908333648 | 0.000338 |
| MGP | -2.883346003 | 5.85E-19 |
| KDR | -2.856988802 | 3.51E-24 |
| ALDH1L2 | -2.749255243 | 0.000597 |
| TGM2 | -2.742967901 | 4.79E-17 |
| DRD2 | -2.707864354 | 0.007043 |
| ALOX5AP | -2.678088804 | 6.89E-06 |
| TAGLN | -2.67413198 | 5.02E-34 |
| COL8A1 | -2.544919939 | 1.59E-12 |
| SELENOP | -2.329975942 | 0.035668 |
| TOX2 | -2.313804905 | 0.035591 |
| WFDC2 | -2.279567504 | 0.000102 |
| MIR137HG | -2.142940565 | 0.038896 |
| SLC7A7 | -2.087552302 | 0.000065 |
| ANPEP | -1.964638827 | 0.003945 |
| C3 | -1.951676084 | 1.11E-30 |
| PTPRB | -1.933491109 | 0.004293 |
| KRT8P3 | -1.915283426 | 0.003079 |
| SLCO2A1 | -1.886321739 | 0.001332 |
| CSF1 | -1.883593433 | 2.39E-12 |
| IL7R | -1.844809704 | 0.010826 |
| AL596244.1 | -1.831755382 | 0.024371 |
| ACSL5 | -1.790662153 | 9.52E-08 |
| SERPINA5 | -1.772609466 | 1.13E-09 |
| CPED1 | -1.746016734 | 0.015922 |
| LBH | -1.741870067 | 1.38E-09 |
| AC037198.1 | -1.729017644 | 0.012144 |
| L1CAM | -1.679768398 | 0.010613 |
| THY1 | -1.659026176 | 0.044468 |
| NNMT | -1.649341609 | 1.55E-09 |
| CDH2 | -1.649154992 | 0.01096 |
| ALPP | -1.614481474 | 0.004953 |
| SDK1 | -1.605917229 | 0.001696 |
| ZNF618 | -1.604356317 | 0.039485 |
| MAPK4 | -1.564725687 | 0.023727 |
| NWD1 | -1.459893665 | 0.002907 |
| AC022075.1 | -1.45856414 | 0.000193 |
| MYADM | -1.438371434 | 0.042316 |
| ADAMTS15 | -1.431838552 | 1.39E-11 |
| THBS1 | -1.422134141 | 3.43E-09 |
| PRR5L | -1.404206504 | 1.58E-06 |
| SAMD4A | -1.397480917 | 2.17E-06 |
| ROR1 | -1.387820804 | 0.002721 |
| SDC3 | -1.372713841 | 0.003185 |
| VTCN1 | -1.359982105 | 0.001494 |
| FILIP1L | -1.335222497 | 0.000087 |
| ZNF488 | -1.327292453 | 0.010069 |
| KRT8 | -1.323255065 | 6.13E-19 |
| KRT81 | -1.322224281 | 0.000168 |
| GABRP | -1.313668399 | 7.07E-07 |
| HEG1 | -1.313321084 | 0.000229 |
| HOXC10 | -1.313103877 | 0.014307 |
| SMOC1 | -1.309430018 | 2.17E-06 |
| IGFBP7 | -1.304973036 | 0.000718 |
| EMP3 | -1.303923506 | 0.007216 |
| FAT4 | -1.302487568 | 5.15E-05 |
| ENC1 | -1.292776496 | 2.15E-05 |
| ANKRD52 | -1.265639539 | 0.04668 |
| GLIPR2 | -1.263008228 | 0.000647 |
| AC026477.1 | -1.259791815 | 0.040886 |
| AC013652.1 | -1.253158388 | 0.014324 |
| MMP2 | -1.220715305 | 0.006901 |
| FN1 | -1.195957847 | 0.020171 |
| FGF2 | -1.19081551 | 0.016121 |
| PLSCR4 | -1.190179602 | 4.08E-07 |
| GAS6-DT | -1.183802812 | 0.039615 |
| RUNX1 | -1.183212386 | 0.015105 |
| IFITM3 | -1.180862094 | 0.00091 |
| TET1 | -1.178169019 | 0.029284 |
| TCEAL3 | -1.166730786 | 2.63E-07 |
| KRT75 | -1.166650815 | 6.85E-05 |
| PMEPA1 | -1.165961913 | 7.3E-06 |
| GDPD5 | -1.164034248 | 6.47E-06 |
| TMEM158 | -1.125176487 | 0.009017 |
| ABLIM3 | -1.120619261 | 0.021011 |
| MARCKSL1 | -1.107663282 | 0.000765 |
| ATF5 | -1.099451058 | 0.01899 |
| PTGER2 | -1.098484809 | 0.000379 |
| LHFPL6 | -1.088761495 | 2.55E-05 |
| CCL28 | -1.087924609 | 0.000773 |
| SULT1E1 | -1.086087606 | 2.37E-05 |
| PTMS | -1.083546725 | 0.013075 |
| LPIN1 | -1.082482828 | 9.21E-09 |
| AL596087.1 | -1.076688307 | 0.022065 |
| COL4A1 | -1.069625851 | 0.004953 |
| GCNT1 | -1.067573891 | 8.54E-08 |
| TNC | -1.065695517 | 0.007285 |
| SLC38A7 | -1.057198956 | 0.034994 |
| ADAM19 | -1.053027085 | 0.041382 |
| B4GALT1 | -1.042534774 | 0.012957 |
| ZFHX3 | -1.024609852 | 0.045978 |
| HOXC13 | -1.020731516 | 0.001311 |
| PXDN | -1.001976884 | 0.000226 |
