## Supplemental Table 2 for "The splicing factor kinase, SR protein kinase 1 (SRPK1) is essential for late events in the human papillomavirus life cycle"

**Supplementary Table 2.** Genes where alternative splicing is significantly altered by SRPIN340 treatment.

| Gene | P value |
| --- | --- |
| MCM10 | 6.16E-29 |
| EIF4G2 | 6.16E-29 |
| SRSF6 | 6.16E-29 |
| SDHB | 6.16E-29 |
| BCAS2 | 6.16E-29 |
| RPLP2 | 6.16E-29 |
| AAAS | 6.16E-29 |
| ALG5 | 6.16E-29 |
| GPSM2 | 6.16E-29 |
| UCK2 | 6.16E-29 |
| P4HA1 | 6.16E-29 |
| ATP6V1D | 6.16E-29 |
| CIRBP | 6.16E-29 |
| KXD1 | 6.16E-29 |
| BLVRB | 6.16E-29 |
| NOP58 | 6.16E-29 |
| PTTG1IP | 6.16E-29 |
| ANXA3 | 6.16E-29 |
| CTNNA1 | 6.16E-29 |
| THBS2 | 6.16E-29 |
| PRKDC | 6.16E-29 |
| COPB1 | 6.16E-29 |
| KRT5 | 6.16E-29 |
| EIF2A | 6.16E-29 |
| TBL1XR1 | 6.16E-29 |
| MCM4 | 6.16E-29 |
| PRXL2A | 6.16E-29 |
| CLNS1A | 6.16E-29 |
| NBR1 | 6.16E-29 |
| CSE1L | 6.16E-29 |
| UBA3 | 6.16E-29 |
| COL7A1 | 6.16E-29 |
| NDUFB9 | 6.16E-29 |
| SCNN1A | 6.16E-29 |
| PCNP | 6.16E-29 |
| TRIM38 | 0.000172 |
| PGLYRP4 | 0.000213 |
| AQP5 | 0.000287 |
| WDR20 | 0.000314 |
| SLC39A9 | 0.000348 |
| ZNF7 | 0.00037 |
| CENPV | 0.000387 |
| FLG | 0.000415 |
| AL589743.1 | 0.000416 |
| TSC1 | 0.000447 |
| FGF1 | 0.000585 |
| SERPINA5 | 0.000658 |
| CCDC120 | 0.000701 |
| DMAP1 | 0.00075 |
| AC105460.1 | 0.000875 |
| ZMAT3 | 0.000903 |
| MYL7 | 0.001055 |
| CYLD | 0.001058 |
| AL162413.1 | 0.001229 |
| MACC1 | 0.001288 |
| CYP1A1 | 0.001446 |
| SUPT7L | 0.001473 |
| FAM133B | 0.001473 |
| OSBPL9 | 0.001473 |
| PDIA4 | 0.001473 |
| CNN2 | 0.001476 |
| NFE2L1 | 0.001483 |
| SMC2 | 0.001723 |
| LAMA3 | 0.001732 |
| ZNF35 | 0.001732 |
| CASP8 | 0.001732 |
| TRIQK | 0.001821 |
| TCP11L2 | 0.001826 |
| N4BP2 | 0.001885 |
| ARTN | 0.00203 |
| NDUFS1 | 0.002111 |
| MAPKAP1 | 0.002244 |
| TRIP12 | 0.002254 |
| AC005476.2 | 0.002485 |
| CD99L2 | 0.002704 |
| KCNK7 | 0.002759 |
| MAB21L4 | 0.002759 |
| FAM185A | 0.002795 |
| AC046134.2 | 0.002954 |
| SPDL1 | 0.003001 |
| GPALPP1 | 0.003126 |
| CHCHD7 | 0.003126 |
| FAM135A | 0.003126 |
| SNHG4 | 0.003126 |
| GPN3 | 0.003126 |
| PDCD10 | 0.003126 |
| UBR3 | 0.003126 |
| SUN1 | 0.003126 |
| SP100 | 0.003249 |
| FANCA | 0.00325 |
| STX2 | 0.003407 |
| ARID1A | 0.003442 |
| LSR | 0.003552 |
| B3GNT9 | 0.003589 |
| TEFM | 0.003602 |
| ZFP64 | 0.003664 |
| PHTF2 | 0.003669 |
| MUC15 | 0.003858 |
| MTRR | 0.003884 |
| TP53AIP1 | 0.003898 |
| FKTN | 0.004025 |
| SMC4 | 0.00409 |
| UBE3D | 0.00409 |
| GUSB | 0.00412 |
| PGAP2 | 0.004147 |
| TMEM134 | 0.004181 |
| IP6K3 | 0.004296 |
| LBR | 0.004365 |
| TGDS | 0.004387 |
| ACSL1 | 0.004419 |
| RPS27A | 0.00442 |
| COQ8B | 0.004541 |
| EPB41 | 0.004541 |
| CTNNAL1 | 0.004592 |
| TXN | 0.004706 |
| IMPDH1 | 0.004729 |
| SLC39A2 | 0.004737 |
| CTSL | 0.004741 |
| PAICS | 0.004752 |
| ZNF557 | 0.004779 |
| CDK18 | 0.004804 |
| NDUFAF2 | 0.004813 |
| METTL23 | 0.004813 |
| SCMH1 | 0.004813 |
| STYXL1 | 0.004813 |
| PAFAH1B3 | 0.004834 |
| FRG1CP | 0.004891 |
| MAP3K7 | 0.004945 |
| TCERG1 | 0.004946 |
| STX18-AS1 | 0.004967 |
| SETD9 | 0.005042 |
| AP1G2 | 0.005063 |
| ACSL5 | 0.005212 |
| TC2N | 0.005249 |
| TNFSF10 | 0.005289 |
| QKI | 0.005507 |
| FBXW7 | 0.005551 |
| CIC | 0.005617 |
| USP54 | 0.005618 |
| SLC38A7 | 0.005672 |
| ZNF2 | 0.005672 |
| TCF25 | 0.005705 |
| CRNN | 0.00582 |
| TRNAU1AP | 0.005838 |
| CDK5RAP1 | 0.005865 |
| LINC00052 | 0.005866 |
| ISG20L2 | 0.005925 |
| PPIP5K1 | 0.005937 |
| VPS54 | 0.006009 |
| CMC1 | 0.006051 |
| SMARCC2 | 0.006054 |
| ANXA8 | 0.006094 |
| DMKN | 0.006133 |
| TREX2 | 0.006204 |
| GABPB1-AS1 | 0.006205 |
| CHAF1A | 0.006476 |
| MIR205HG | 0.006687 |
| TUG1 | 0.006694 |
| PIF1 | 0.006715 |
| FBH1 | 0.006715 |
| ODF2 | 0.006734 |
| SGK3 | 0.006738 |
| LAMTOR3 | 0.006763 |
| TIA1 | 0.006797 |
| RACGAP1 | 0.006811 |
| MPDU1 | 0.006827 |
| FNIP1 | 0.006853 |
| HSPBP1 | 0.006853 |
| SNHG11 | 0.007063 |
| DPH5 | 0.007091 |
| PSRC1 | 0.007148 |
| AP3S2 | 0.007331 |
| OCEL1 | 0.007334 |
| ZNF462 | 0.007336 |
| CCBE1 | 0.007366 |
| ODR4 | 0.00742 |
| PPIE | 0.00747 |
| SEC11A | 0.007477 |
| CRYAB | 0.007683 |
| GMIP | 0.007707 |
| CTSC | 0.007763 |
| ZNF214 | 0.007763 |
| RPF2 | 0.007771 |
| ARMC6 | 0.007771 |
| CSNK1G3 | 0.0078 |
| TAF12 | 0.007958 |
| OPLAH | 0.007978 |
| SNX25 | 0.008048 |
| STK38L | 0.00805 |
| SYCP2 | 0.00807 |
| RANBP3 | 0.008303 |
| ATXN7L3 | 0.008303 |
| SHPRH | 0.008388 |
| ATP2C1 | 0.008477 |
| ZNF75D | 0.008508 |
| FAM111A | 0.00851 |
| RNPEP | 0.008573 |
| TRIM29 | 0.008596 |
| VEGFB | 0.008671 |
| PHF3 | 0.008731 |
| HMGA1 | 0.008731 |
| PAX8-AS1 | 0.009041 |
| HINFP | 0.009057 |
| SP110 | 0.009133 |
| ZNF691 | 0.009133 |
| BAIAP2 | 0.009133 |
| TP73-AS1 | 0.009248 |
| SLC38A9 | 0.00959 |
| SLC22A15 | 0.00964 |
| HERC1 | 0.009711 |
| FAHD2A | 0.009795 |
| FAM53C | 0.009852 |
| LCORL | 0.009852 |
| EIF4G1 | 0.009852 |
| KLK13 | 0.009964 |
| FAM120A | 0.009964 |
| RAC3 | 0.009964 |
| SEPTIN2 | 0.009964 |
| PDE6D | 0.010197 |
| PLEC | 0.010287 |
| TPM1 | 0.01033 |
| CYP2U1 | 0.010394 |
| TERF1 | 0.010399 |
| ZNF12 | 0.010452 |
| CARD8 | 0.01049 |
| DCAF11 | 0.010579 |
| NOP14-AS1 | 0.010664 |
| DCAF16 | 0.0107 |
| TFB1M | 0.0107 |
| BRCA1 | 0.010822 |
| LIX1L | 0.011015 |
| TMEM63A | 0.011071 |
| ZNHIT1 | 0.011098 |
| ZNF236 | 0.011258 |
| FUZ | 0.011258 |
| PDZK1 | 0.011286 |
| TCAF1 | 0.011521 |
| KIF2C | 0.011526 |
| PRELID3B | 0.011566 |
| SMUG1 | 0.011705 |
| STARD4 | 0.011733 |
| GABRP | 0.011741 |
| WDR66 | 0.011775 |
| TMEM164 | 0.011775 |
| TSPAN17 | 0.011835 |
| SNRPN | 0.011857 |
| HNRNPDL | 0.011936 |
| MGA | 0.011985 |
| MSTO1 | 0.012018 |
| PDPR | 0.012384 |
| DDIT4 | 0.012439 |
| MGRN1 | 0.012471 |
| SLC29A1 | 0.012645 |
| KHDRBS1 | 0.012757 |
| PORCN | 0.012822 |
| HNRNPR | 0.012863 |
| CNOT6L | 0.012944 |
| ATPSCKMT | 0.012976 |
| RGS10 | 0.013022 |
| NEIL1 | 0.013045 |
| COPE | 0.013129 |
| ANKRD11 | 0.013151 |
| FDPS | 0.013172 |
| INTS2 | 0.013205 |
| MED8 | 0.013236 |
| PPME1 | 0.013236 |
| PRKRA | 0.013236 |
| NPHP3 | 0.013236 |
| RMDN1 | 0.013236 |
| MAN2C1 | 0.013236 |
| SH3GLB2 | 0.013236 |
| ZDHHC17 | 0.013236 |
| MPZL1 | 0.013236 |
| TIMMDC1 | 0.013236 |
| ZNF639 | 0.013236 |
| CLEC2D | 0.013236 |
| POMK | 0.013236 |
| MPV17 | 0.013236 |
| USO1 | 0.013236 |
| SNRK | 0.013236 |
| C11orf74 | 0.013236 |
| PSMD3 | 0.013245 |
| NCOR1 | 0.013245 |
| WWP2 | 0.013322 |
| GMPR2 | 0.013322 |
| CRYBG2 | 0.013399 |
| PC | 0.013434 |
| PTPRF | 0.01356 |
| ZNF121 | 0.013601 |
| IRX2 | 0.013646 |
| MTX2 | 0.01369 |
| PATZ1 | 0.013746 |
| POLL | 0.013949 |
| JMY | 0.013949 |
| RHBDD3 | 0.013952 |
| ZC3H7B | 0.013982 |
| DDIAS | 0.014051 |
| KLK12 | 0.014075 |
| TWNK | 0.014111 |
| TM2D1 | 0.014262 |
| DEXI | 0.014336 |
| ZNF638 | 0.01434 |
| HMGN1 | 0.014454 |
| GFM1 | 0.014575 |
| OSMR | 0.014711 |
| GALNT12 | 0.014935 |
| P2RX4 | 0.014953 |
| EIF2B4 | 0.015268 |
| FKBP7 | 0.015268 |
| LGALS7B | 0.015268 |
| MTERF4 | 0.015268 |
| USP28 | 0.015268 |
| TMEM175 | 0.015268 |
| PRKRIP1 | 0.015268 |
| NCBP3 | 0.015268 |
| NBPF26 | 0.015268 |
| GRHL1 | 0.015287 |
| TTC3 | 0.015318 |
| SMIM19 | 0.015327 |
| TMEM267 | 0.015433 |
| AC006001.3 | 0.015443 |
| TXNL4A | 0.015515 |
| GOLGA1 | 0.015515 |
| TCEAL9 | 0.015523 |
| LRATD2 | 0.015531 |
| FAM114A2 | 0.015618 |
| SSBP2 | 0.015658 |
| NLRC5 | 0.015658 |
| RAB28 | 0.015734 |
| OXR1 | 0.015821 |
| ERCC3 | 0.015902 |
| HNRNPH1 | 0.015902 |
| SRD5A3 | 0.015977 |
| ZNF500 | 0.016001 |
| RDH16 | 0.016011 |
| SEC62 | 0.016011 |
| UCHL5 | 0.016011 |
| RUNDC3A | 0.016029 |
| MARCHF5 | 0.016065 |
| GBF1 | 0.016173 |
| GLYR1 | 0.016355 |
| ITGA6 | 0.016404 |
| ANK3 | 0.016472 |
| LCOR | 0.016504 |
| PIKFYVE | 0.016509 |
| ARAP3 | 0.01663 |
| PI4K2B | 0.01663 |
| DAG1 | 0.01664 |
| UBE2D3 | 0.016736 |
| SCYL3 | 0.016796 |
| MTMR6 | 0.016856 |
| CAMK2D | 0.016856 |
| HELZ | 0.016858 |
| STIL | 0.01696 |
| PRPF39 | 0.017 |
| EEF1AKMT2 | 0.017021 |
| WARS1 | 0.017128 |
| THOC2 | 0.017131 |
| TP53I3 | 0.017215 |
| PHC3 | 0.017442 |
| ZNF263 | 0.017564 |
| AP1G1 | 0.017565 |
| ASAP1 | 0.017603 |
| MINDY1 | 0.017632 |
| GLYCTK | 0.017686 |
| CBS | 0.017746 |
| ZNF33A | 0.017797 |
| MBD1 | 0.017915 |
| ANKRD35 | 0.017915 |
| MOCS2 | 0.018092 |
| TDG | 0.018092 |
| LINC02434 | 0.018408 |
| LCMT1 | 0.018509 |
| DHX30 | 0.01863 |
| SLC11A2 | 0.018727 |
| HPS4 | 0.018742 |
| SLTM | 0.018746 |
| CPSF2 | 0.018777 |
| DDB2 | 0.018844 |
| HACL1 | 0.019211 |
| CUTC | 0.019223 |
| NBEAL2 | 0.019442 |
| RIC8B | 0.019459 |
| DDX49 | 0.01946 |
| NFYC | 0.019504 |
| ZDHHC13 | 0.0196 |
| CNTNAP3 | 0.01962 |
| ZNF613 | 0.019751 |
| BIRC2 | 0.019804 |
| NPEPPS | 0.019804 |
| IFT57 | 0.019804 |
| CDK5RAP2 | 0.019804 |
| NAT9 | 0.019804 |
| ATP9B | 0.019804 |
| LUC7L | 0.019804 |
| ZC3H18 | 0.019804 |
| CDC25C | 0.019804 |
| LRBA | 0.019804 |
| GAS5 | 0.019804 |
| LARP4 | 0.019804 |
| ZNF451 | 0.019887 |
| ARL6IP4 | 0.019887 |
| ZNF620 | 0.01993 |
| NFYA | 0.020061 |
| MST1R | 0.020066 |
| ZWILCH | 0.020092 |
| EXD2 | 0.020092 |
| GSTP1 | 0.020143 |
| MAPK8 | 0.020169 |
| ARRDC1 | 0.020169 |
| PRR15L | 0.020204 |
| TACC3 | 0.020204 |
| AC012085.1 | 0.020204 |
| TNNT1 | 0.020204 |
| RPL38 | 0.020204 |
| MBNL1 | 0.020204 |
| PGS1 | 0.020252 |
| TMEM241 | 0.020254 |
| MARK2 | 0.020254 |
| NBPF8 | 0.020274 |
| SEMA4F | 0.020335 |
| ADHFE1 | 0.020394 |
| NSD2 | 0.020467 |
| PMS2 | 0.02071 |
| SLC26A2 | 0.020976 |
| CCNL1 | 0.020998 |
| NIT2 | 0.021019 |
| MAP2 | 0.021028 |
| ANAPC4 | 0.02109 |
| LTBP4 | 0.021101 |
| DENND2B | 0.021135 |
| NAP1L4 | 0.021302 |
| CTNND1 | 0.021312 |
| SLC27A2 | 0.021339 |
| NNT | 0.021402 |
| U2AF1L4 | 0.021405 |
| ARIH2 | 0.021463 |
| IL18 | 0.021561 |
| ZNF207 | 0.021596 |
| ZC3H4 | 0.021605 |
| ZBTB8OS | 0.021642 |
| IQGAP3 | 0.021651 |
| PCBP2 | 0.021724 |
| HOTAIR | 0.021724 |
| ARHGAP11A | 0.021728 |
| NAV2 | 0.021762 |
| FHL2 | 0.02179 |
| UTRN | 0.021894 |
| ZUP1 | 0.021912 |
| ZNF81 | 0.022032 |
| CYB561 | 0.022142 |
| PRMT1 | 0.022202 |
| P4HA2 | 0.022248 |
| PRPF40B | 0.022273 |
| CBWD1 | 0.02233 |
| METTL1 | 0.02237 |
| MATN2 | 0.022551 |
| IDH3A | 0.022554 |
| CSDE1 | 0.022671 |
| DPY19L4 | 0.022792 |
| AGAP3 | 0.02286 |
| LINC02541 | 0.02286 |
| RNF41 | 0.022969 |
| JADE2 | 0.023093 |
| HNRNPC | 0.02312 |
| SNX14 | 0.02317 |
| ARRDC3 | 0.02317 |
| AK2 | 0.023175 |
| FIBP | 0.023214 |
| EIF4A2 | 0.023339 |
| SNRPG | 0.02337 |
| SERPINF1 | 0.023376 |
| STAG2 | 0.02351 |
| SLC19A1 | 0.023544 |
| TRIM5 | 0.023544 |
| ALG10B | 0.023561 |
| TEAD2 | 0.023689 |
| DUOXA1 | 0.023691 |
| BX890604.2 | 0.023792 |
| USP8 | 0.023894 |
| TRAPPC2L | 0.023907 |
| DGKA | 0.023914 |
| HSPB11 | 0.02411 |
| PRSS23 | 0.02411 |
| DSG3 | 0.02411 |
| HAGHL | 0.02411 |
| PARP8 | 0.02411 |
| CEP164 | 0.02411 |
| KRT13 | 0.02411 |
| TLK2 | 0.02411 |
| TCFL5 | 0.02411 |
| BAZ2B | 0.02411 |
| MRPL52 | 0.02411 |
| KIFAP3 | 0.02411 |
| MRPL22 | 0.02411 |
| ZNF428 | 0.02411 |
| CDC42SE1 | 0.02411 |
| LYPLA1 | 0.02411 |
| NDC1 | 0.02411 |
| IP6K2 | 0.02411 |
| NAPA | 0.02411 |
| CHD1L | 0.024114 |
| PBRM1 | 0.024146 |
| PDLIM2 | 0.024149 |
| KREMEN2 | 0.024155 |
| TTC13 | 0.024196 |
| POT1 | 0.024348 |
| RNH1 | 0.024384 |
| SS18 | 0.024389 |
| HTATIP2 | 0.02439 |
| GON7 | 0.02446 |
| IFT81 | 0.024473 |
| FBXL12 | 0.024542 |
| ZKSCAN8 | 0.024566 |
| ARSJ | 0.024566 |
| ZBTB24 | 0.024593 |
| SMCHD1 | 0.024623 |
| RDH11 | 0.024662 |
| HOMER2 | 0.024662 |
| ZNF266 | 0.024695 |
| ARHGAP10 | 0.024723 |
| ZYG11A | 0.024817 |
| HAGH | 0.024817 |
| ZNF3 | 0.024817 |
| MARCHF7 | 0.024817 |
| MMS19 | 0.024817 |
| GALE | 0.024817 |
| CSNK1E | 0.024817 |
| PSPC1 | 0.024817 |
| ELL2 | 0.024817 |
| KANK4 | 0.024836 |
| FAP | 0.025042 |
| KCTD20 | 0.025273 |
| SKI | 0.025303 |
| CHD7 | 0.025303 |
| ZSCAN20 | 0.025303 |
| WDYHV1 | 0.025303 |
| ELK3 | 0.025303 |
| ASAH2B | 0.025327 |
| RARS2 | 0.025492 |
| PTPRG-AS1 | 0.025532 |
| ZBED5 | 0.025755 |
| RUFY1 | 0.025824 |
| ATP13A2 | 0.026039 |
| KRT8 | 0.026106 |
| TMEM107 | 0.026121 |
| USP31 | 0.026148 |
| C1orf109 | 0.026229 |
| TMEM99 | 0.026268 |
| TMPO | 0.026271 |
| SEC22C | 0.026382 |
| ALAS1 | 0.026413 |
| UBR5 | 0.026522 |
| DRAM2 | 0.026529 |
| ZNF184 | 0.026671 |
| ZFAND6 | 0.026789 |
| OGFOD2 | 0.026809 |
| ARAP1 | 0.026812 |
| LARP1 | 0.026817 |
| MED24 | 0.026877 |
| SRPX2 | 0.026974 |
| MLLT1 | 0.026976 |
| THRAP3 | 0.027086 |
| ZNF146 | 0.027122 |
| ALKBH3 | 0.027122 |
| MT1L | 0.027122 |
| MIGA2 | 0.027149 |
| DSN1 | 0.027149 |
| STAT3 | 0.027196 |
| RHBDD1 | 0.027373 |
| PARK7 | 0.027478 |
| AC093752.1 | 0.027528 |
| KIFC3 | 0.027591 |
| BIRC5 | 0.027619 |
| SLC35F6 | 0.027845 |
| RPAIN | 0.027987 |
| MOV10 | 0.028078 |
| SREBF1 | 0.028143 |
| IFT80 | 0.02821 |
| DCBLD1 | 0.028391 |
| POMT1 | 0.02842 |
| OAT | 0.02842 |
| NKIRAS2 | 0.028481 |
| PRMT2 | 0.028582 |
| SLC25A29 | 0.028596 |
| SCARB1 | 0.028596 |
| KCTD3 | 0.028596 |
| KIF23 | 0.028596 |
| RPS14 | 0.028596 |
| AC091230.1 | 0.028596 |
| RAD17 | 0.028596 |
| DDB1 | 0.028596 |
| ZC3H11A | 0.028596 |
| CLK1 | 0.028747 |
| DECR1 | 0.028842 |
| FBXO7 | 0.028868 |
| CMTM4 | 0.028887 |
| MFSD14B | 0.028978 |
| DENND2C | 0.028978 |
| SNUPN | 0.028998 |
| KRTDAP | 0.029045 |
| PIGW | 0.029164 |
| POMT2 | 0.029216 |
| SLC12A9 | 0.029365 |
| RPL7L1 | 0.029403 |
| FECH | 0.029454 |
| TRA2A | 0.029454 |
| AP3S1 | 0.02947 |
| IMMP1L | 0.029558 |
| G2E3 | 0.029564 |
| CPNE7 | 0.029759 |
| DGKD | 0.029759 |
| CASC19 | 0.029831 |
| PMEPA1 | 0.029913 |
| ABCG1 | 0.030044 |
| NQO1 | 0.030232 |
| MPRIP | 0.03038 |
| ESCO1 | 0.030437 |
| FBXL2 | 0.030477 |
| ITFG2 | 0.030483 |
| PTPRE | 0.030486 |
| STAM | 0.030557 |
| ACACA | 0.03078 |
| OSBPL1A | 0.03078 |
| SNHG17 | 0.030811 |
| RHOT1 | 0.030817 |
| ANXA2 | 0.0309 |
| PTGES2 | 0.030906 |
| RIOK3 | 0.030931 |
| CEP290 | 0.030956 |
| ZC2HC1A | 0.030997 |
| TUBGCP4 | 0.031087 |
| HP1BP3 | 0.031441 |
| RC3H2 | 0.031486 |
| PLRG1 | 0.031504 |
| CHEK1 | 0.031504 |
| CCDC77 | 0.031557 |
| KRIT1 | 0.03157 |
| AKR1A1 | 0.031663 |
| PRKAB1 | 0.031909 |
| SRRM1 | 0.032007 |
| SHARPIN | 0.03209 |
| TBL2 | 0.032199 |
| MINDY3 | 0.032199 |
| TXNRD1 | 0.032199 |
| RBM41 | 0.032199 |
| UBQLN1 | 0.032299 |
| ETHE1 | 0.032328 |
| LRRC42 | 0.032328 |
| MCFD2 | 0.032408 |
| SDR39U1 | 0.032415 |
| OCIAD1 | 0.032573 |
| UBAC2 | 0.032573 |
| ABCC10 | 0.032672 |
| NT5C3A | 0.032672 |
| ZDHHC6 | 0.032672 |
| CDK10 | 0.032672 |
| ACTR1B | 0.032678 |
| UBE3A | 0.032736 |
| AL365205.1 | 0.032759 |
| USP47 | 0.032811 |
| MAD2L1 | 0.032889 |
| RECQL | 0.032996 |
| KRI1 | 0.033083 |
| ZNF532 | 0.033255 |
| FAM111B | 0.033337 |
| RHOT2 | 0.033455 |
| MCM3 | 0.033459 |
| MYO9B | 0.033472 |
| MAX | 0.033472 |
| ILKAP | 0.033608 |
| RNPC3 | 0.033623 |
| ENOPH1 | 0.033693 |
| RRNAD1 | 0.033696 |
| LAMB3 | 0.033908 |
| MED17 | 0.033908 |
| TANGO2 | 0.033908 |
| PRUNE1 | 0.033915 |
| TMEM131L | 0.033955 |
| SAT1 | 0.033997 |
| C18orf25 | 0.034153 |
| KLK11 | 0.034163 |
| TXLNA | 0.034177 |
| PFDN5 | 0.034219 |
| KLHL24 | 0.034343 |
| CA12 | 0.03446 |
| DNPEP | 0.034465 |
| AC093157.1 | 0.034567 |
| CINP | 0.034578 |
| HOOK2 | 0.034578 |
| DUSP22 | 0.034608 |
| GSTO2 | 0.034674 |
| MORF4L1 | 0.034708 |
| ZDHHC20 | 0.034805 |
| CDH16 | 0.03494 |
| LRFN4 | 0.03499 |
| RBM10 | 0.035047 |
| FXYD3 | 0.035059 |
| CAMTA1 | 0.035099 |
| RPL22L1 | 0.035099 |
| NEMF | 0.035099 |
| SLC37A3 | 0.035099 |
| PLAA | 0.035099 |
| MTFR1L | 0.035099 |
| FADS3 | 0.035099 |
| UBXN1 | 0.035099 |
| TBK1 | 0.035099 |
| C1orf43 | 0.035099 |
| TENM2 | 0.035099 |
| GRAMD2A | 0.035131 |
| SF3A3 | 0.035243 |
| TRMU | 0.0354 |
| SULT1E1 | 0.035426 |
| MXD4 | 0.035518 |
| DALRD3 | 0.035592 |
| MVP | 0.035637 |
| BSG | 0.035637 |
| FBXO38 | 0.035653 |
| WBP2 | 0.035653 |
| WDR75 | 0.035653 |
| CAST | 0.035653 |
| WASHC4 | 0.035653 |
| BLVRA | 0.035653 |
| ABHD18 | 0.035653 |
| CDK2AP1 | 0.035653 |
| GPATCH8 | 0.035653 |
| OSBPL8 | 0.035653 |
| RPARP-AS1 | 0.035653 |
| ZFP90 | 0.035683 |
| GABPB1 | 0.035732 |
| TRAPPC10 | 0.035747 |
| SUGP2 | 0.035787 |
| MAGI1 | 0.03579 |
| BCS1L | 0.035864 |
| CBSL | 0.035894 |
| GANC | 0.035991 |
| VRK2 | 0.03616 |
| ADD3 | 0.036424 |
| PNO1 | 0.036641 |
| NISCH | 0.036641 |
| AKR1E2 | 0.036668 |
| TP73 | 0.036693 |
| CORO6 | 0.037 |
| PPIL2 | 0.037006 |
| PRDM15 | 0.037086 |
| STRN3 | 0.037086 |
| PACRGL | 0.037146 |
| HTATSF1 | 0.037206 |
| FEZ2 | 0.03722 |
| NUDT12 | 0.037271 |
| FANCI | 0.037372 |
| MOGS | 0.037403 |
| PTPRZ1 | 0.037605 |
| FRMD4B | 0.037697 |
| AAMDC | 0.03771 |
| BECN1 | 0.037725 |
| ENY2 | 0.03775 |
| CD27-AS1 | 0.03775 |
| ZBTB38 | 0.03775 |
| DHRS4 | 0.03775 |
| SNHG14 | 0.03775 |
| LMNB2 | 0.03775 |
| CKAP2L | 0.03775 |
| PTPN3 | 0.03775 |
| TMEM161B | 0.03775 |
| SLC35B1 | 0.03775 |
| KCNJ15 | 0.03775 |
| ZNF283 | 0.03775 |
| SLC35A3 | 0.03775 |
| CNTROB | 0.03775 |
| RIN2 | 0.03775 |
| CAPN10 | 0.03775 |
| DERL2 | 0.03775 |
| COMMD3 | 0.03775 |
| SEC23A | 0.03775 |
| DNAJC7 | 0.03775 |
| TOP2B | 0.03775 |
| CDC42SE2 | 0.03775 |
| SPICE1 | 0.03775 |
| LPIN3 | 0.03775 |
| AVL9 | 0.03775 |
| NUDT2 | 0.03775 |
| BPNT1 | 0.03775 |
| NFRKB | 0.03775 |
| KDM5D | 0.03775 |
| RAB11FIP2 | 0.03775 |
| POMGNT1 | 0.03775 |
| KLF6 | 0.03775 |
| PSMA4 | 0.03775 |
| BLNK | 0.037945 |
| BRD3 | 0.038009 |
| ACAA1 | 0.038074 |
| IL17RE | 0.038102 |
| ACIN1 | 0.038212 |
| FAM219A | 0.038344 |
| TMCO4 | 0.03838 |
| ANO10 | 0.038472 |
| THUMPD3-AS1 | 0.038491 |
| EXTL3 | 0.038561 |
| FANK1 | 0.038561 |
| EEF2KMT | 0.038561 |
| JKAMP | 0.038561 |
| ZNF76 | 0.03863 |
| SERPINB13 | 0.038639 |
| FANCD2 | 0.038672 |
| ROBO1 | 0.038674 |
| PRSS27 | 0.038692 |
| PNPLA4 | 0.038692 |
| PAM | 0.038786 |
| STAG3L5P-PVRIG2P-PILRB | 0.038969 |
| CSNK2A1 | 0.039012 |
| POLD4 | 0.039012 |
| PPP4R2 | 0.039012 |
| SLC25A36 | 0.039073 |
| PPP2R1B | 0.039109 |
| RAD51D | 0.039256 |
| CLASP2 | 0.039333 |
| CNOT8 | 0.039409 |
| TRRAP | 0.039483 |
| BZW2 | 0.039568 |
| AP4E1 | 0.039778 |
| NME6 | 0.039852 |
| ERI3 | 0.039872 |
| S100A2 | 0.039872 |
| POLQ | 0.039872 |
| TAF2 | 0.040046 |
| LIMA1 | 0.040054 |
| ACADVL | 0.040054 |
| RNF8 | 0.040054 |
| NUP98 | 0.040054 |
| ZNF860 | 0.040061 |
| LINC01322 | 0.040138 |
| B3GALNT2 | 0.040168 |
| MRPL24 | 0.040298 |
| PLSCR1 | 0.040318 |
| PIGG | 0.040408 |
| SRRD | 0.040431 |
| OSBPL3 | 0.040496 |
| ACAD10 | 0.040658 |
| TBC1D5 | 0.04066 |
| AC016876.2 | 0.040711 |
| EI24 | 0.040753 |
| KIF18B | 0.040833 |
| ZNF84 | 0.041161 |
| STARD10 | 0.041169 |
| NSUN6 | 0.041379 |
| JUP | 0.041441 |
| PPP4C | 0.041489 |
| SYPL1 | 0.041525 |
| TTLL5 | 0.041678 |
| TENT2 | 0.041695 |
| PSME4 | 0.041739 |
| GPR89B | 0.041769 |
| PPIH | 0.041915 |
| RANBP1 | 0.041915 |
| NPRL2 | 0.04211 |
| KANSL2 | 0.042153 |
| CGRRF1 | 0.042153 |
| FBXO22 | 0.042159 |
| CEP44 | 0.042172 |
| GTF2I | 0.042172 |
| SOX13 | 0.042193 |
| MAPK10 | 0.042251 |
| ARMCX6 | 0.0423 |
| NASP | 0.042346 |
| NUP107 | 0.042394 |
| TSEN15 | 0.042414 |
| TMEM62 | 0.042479 |
| STARD5 | 0.0426 |
| TMTC3 | 0.042764 |
| APEH | 0.042865 |
| RFC5 | 0.042865 |
| AFMID | 0.043088 |
| PLEK2 | 0.0431 |
| TMEM138 | 0.043162 |
| PPDPF | 0.043308 |
| PLEKHF1 | 0.043514 |
| ELMOD3 | 0.043592 |
| TRMT10A | 0.043689 |
| USP14 | 0.043692 |
| MFSD13A | 0.043773 |
| ENC1 | 0.043792 |
| PHKB | 0.043817 |
| COA1 | 0.043838 |
| TUSC3 | 0.043878 |
| STAMBP | 0.043881 |
| SAFB | 0.044236 |
| BCCIP | 0.04425 |
| CKAP2 | 0.04434 |
| LZIC | 0.044467 |
| SETDB2 | 0.044467 |
| YME1L1 | 0.044467 |
| SETD5 | 0.044467 |
| TRIO | 0.044505 |
| FBXO25 | 0.044548 |
| PPP6R3 | 0.044691 |
| RIPOR2 | 0.044978 |
| PTAR1 | 0.045024 |
| SCEL | 0.045026 |
| MATR3 | 0.045201 |
| LNCAROD | 0.04546 |
| NCKAP5L | 0.04546 |
| SNX12 | 0.045541 |
| SVBP | 0.04558 |
| AC245041.2 | 0.045652 |
| MAGOHB | 0.045976 |
| EIF3E | 0.04609 |
| WDR92 | 0.046107 |
| KLK10 | 0.046125 |
| FUBP1 | 0.046252 |
| NUDT9 | 0.046389 |
| TIAL1 | 0.046996 |
| BLOC1S6 | 0.047085 |
| ZNF561 | 0.047103 |
| AGTRAP | 0.047179 |
| CLDN12 | 0.047195 |
| FAM214B | 0.047214 |
| APEX1 | 0.047223 |
| COX10-AS1 | 0.047268 |
| USP19 | 0.047362 |
| MALT1 | 0.047421 |
| SMIM22 | 0.047421 |
| MON1B | 0.047421 |
| TALDO1 | 0.047421 |
| AP3D1 | 0.047421 |
| CMSS1 | 0.047421 |
| CENPK | 0.047421 |
| DCTN5 | 0.047421 |
| SART1 | 0.04881 |
| DNTTIP1 | 0.04881 |
| ATRX | 0.04905 |
| FAM200B | 0.049142 |
| ELAC2 | 0.049372 |
| LINC02660 | 0.049405 |
| EIF3B | 0.049779 |
